# Min waves without MinC can pattern FtsA-FtsZ filaments on model membranes

**DOI:** 10.1101/2021.11.15.468671

**Authors:** Elisa Godino, Anne Doerr, Christophe Danelon

## Abstract

Although the essential proteins that drive bacterial cytokinesis have been identified and reconstituted in vitro, the precise mechanisms by which they dynamically interact to enable symmetrical division are largely unknown. In *Escherichia coli*, cell division begins with the formation of a proto-ring composed of FtsZ and its membrane-tethering proteins FtsA and ZipA. In the broadly proposed molecular scenario for ring positioning, Min waves composed of MinD and MinE distribute the FtsZ-polymerization inhibitor MinC away from mid-cell, where the Z-ring can form. Therefore, MinC is believed to be an essential element connecting the Min and FtsZ systems. Here, by using cell-free gene expression on planar lipid membranes, we demonstrate that MinDE drive the formation of dynamic, antiphase patterns of FtsZ-FtsA co-filaments even in the absence of MinC. This behavior is also observed when the proteins are compartmentalized inside microdroplets. These results suggest that Z-ring positioning may be achieved with a more minimal set of proteins than previously envisaged, providing a fresh perspective about the role of MinC. Moreover, we propose that MinDE oscillations may constitute the minimal localization mechanism of an FtsA-FtsZ constricting ring in a prospective synthetic cell.

## Introduction

In *Escherichia coli* bacteria, a ring-forming multiprotein complex drives membrane constriction at the future division site^1^. The proto-ring is composed of the three proteins FtsZ, FtsA and ZipA^2,3^. FtsZ is a tubulin homologue with GTPase activity that can polymerize into protofilaments^4,5^. FtsZ has no affinity for lipids and is anchored to the membrane by ZipA and the actin homologue FtsA^6–8^. This set of proteins acts as a scaffold, recruiting other factors to form a mature divisome. The Min system in *E. coli* is composed of the three proteins MinC, MinD, and MinE. Together, they provide the localization cues that restrict the assembly of the FtsZ ring to the middle of the cell, following a commonly accepted sequence of molecular events^9^. MinD is an ATPase that dimerizes and is recruited on the membrane when bound to ATP. Once at the membrane, MinD interacts with MinE, which stimulates the ATPase activity of MinD leading to the release of both proteins from the membrane. This dynamic interplay between MinD and MinE, together with the geometrical constraints of the cell, cause oscillating gradients of the two proteins at the cell poles^10–13^. MinC travels with the MinDE waves, where it inhibits the assembly of FtsZ protofilaments. As a result of the Min oscillations, the time-averaged concentration of the MinD-MinC complex is higher at the poles, encouraging the formation of the Z-ring to mid-cell^14–16^.

Despite numerous in vitro studies about membrane-anchored FtsZ^17–20^ and Min dynamics^16,21–23^, data about the spatiotemporal regulation of FtsA-FtsZ cytoskeletal structures by MinDEC are scarce^24^. It is known that MinC is required for correct placement of the division site in vivo^10,13^. On the other hand, several reports suggest that MinDE oscillations may act as a spatial regulator of membrane proteins^25–27^. The abundances of the membrane proteome in wild-type and Δ*min E. coli* cells were compared, revealing that Min oscillating gradients modulate protein association with the inner membrane^25^. Moreover, MinDE dynamic patterns spatiotemporally regulate and transport peripheral membrane proteins on supported lipid bilayers (SLBs) and in microcompartments^26,27^. These recent findings raise the question of whether the oscillating MinDE gradients could influence membrane localization of FtsA-FtsZ filaments in the absence of MinC. Interestingly, when FtsZ-YFP-MTS (a chimera of truncated FtsZ, YFP, and the membrane-targeting sequence of MinD) was combined with MinDE proteins on an SLB, static FtsZ-YFP-MTS networks were not affected by the Min surface waves^26^. The use of a chimeric FtsZ in these in vitro assays leaves open questions, since the ability of MinDE oscillating gradients to influence the lateral distribution of FtsZ might be different with the native membrane anchor FtsA.

Herein, we set out to elucidate this question by combining MinDE(C), where ‘(C)’ indicates the presence or absence of MinC, and FtsA-FtsZ in open and closed model membrane systems. As a medium for our cell-free assays, we use the PURE system, a minimal gene expression system reconstituted primarily from *E. coli* constituents^28,29^. The activity of PURE-expressed FtsA and MinDEC from single-gene constructs has already been validated^24,30^. Here, we design a DNA template containing the *E. coli* genes *ftsA*, *minD* and *minE* concatenated in the form of three transcriptional units (tri-cistron). All proteins are expressed at physiologically relevant levels and in active states, allowing us to examine the interplay between the spatial organization of FtsA-FtsZ cytoskeletal structures and MinDE(C) membrane patterns on SLBs and within microdroplets. We find that MinDE surface waves act as a spatial regulator of FtsA-FtsZ filaments in a MinC-independent manner. The possible implications regarding bacterial cell division and implementation of a minimal FtsZ ring in synthetic cells are discussed.

## Results

### Cell-free co-expression of FtsA, MinD and MinE at physiological levels

As an experimental design strategy to co-reconstitute the Min and FtsZ subsystems, we created a multicistronic DNA template containing the genes of the *E. coli* proteins FtsA, MinD, and MinE (Fig. 1 A). The DNA template (5 nM) was constitutively expressed in PURE*frex*2.0 and the abundance of the three synthesized proteins was quantified by targeted liquid chromatography-coupled mass spectrometry (LC-MS) (Fig. 1 A and Fig. S1). After 3 hours of expression, the obtained concentrations were 1.5 ± 0.1 μM for FtsA, 4.2 ± 0.9 μM for MinD and 3.2 ± 0.4 μM for MinE (mean ± SD, three biological replicates) (Fig. 1 B). These values give a good approximation of the total amounts of full-length proteins but may overestimate the concentrations of active proteins. Similar concentration values were reported in vivo: ~0.5 μM for FtsA^31^, and an equimolar amount of MinD and MinE ranging between 1 and 3 μM^32,33^. Kinetics of FtsA, MinD and MinE production revealed that most of FtsA and MinD proteins were synthesized within the first 3 hours of co-expression (Fig. 1 C). The apparent translation rate was calculated by fitting a phenomenological model to the experimental data yielding values of 0.018 ± 0.002 μM min^−1^ for FtsA, 0.043 ± 0.016 μM min^−1^ for MinD and 0.027 ± 0.002 μM min^−1^ for MinE (mean ± SD of fitted parameter values, three biological replicates). The expression lifespan, defined as the time point at which protein production stops, was determined for each protein: 119 ± 28 min (FtsA), 144 ± 28 min (MinD) and 250 ± 16 min (MinE). These values are consistent with those obtained from kinetic profiles of other PURE-expressed proteins^34,35^.

**Figure 1.**
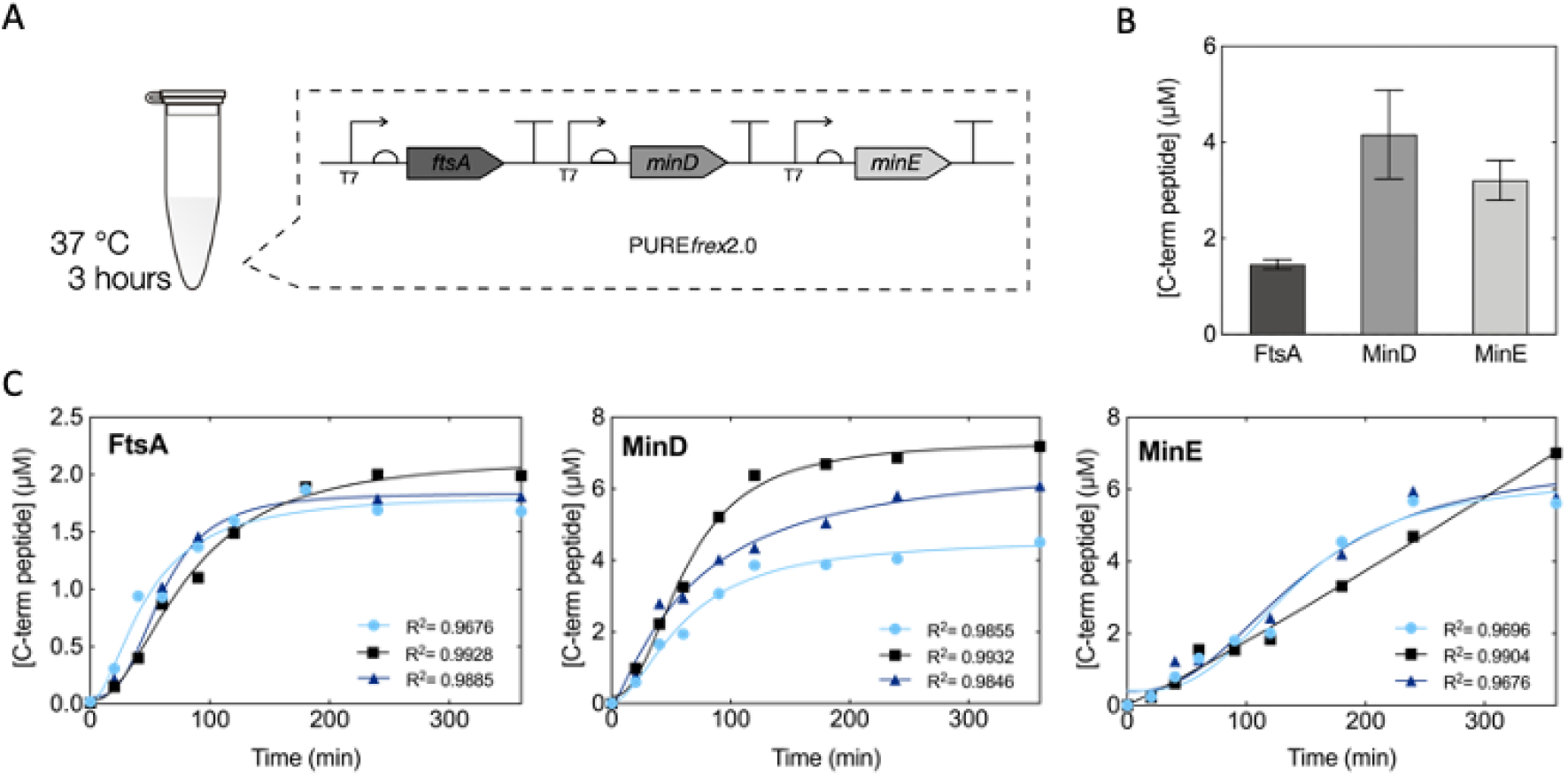
Quantification of cell-free expressed FtsA, MinD and MinE proteins from a single DNA construct. (A) Each gene, *ftsA*, *minD* and *minE*, was placed under its own T7 promoter, ribosome binding site and T7 transcription terminator and was constitutively expressed with PURE*frex*2.0 in test-tube reactions. (B) Quantitative LC-MS analysis of FtsA, MinD and MinE protein production after 3 hours of expression at 37 °C. Abundance of the most C-terminal proteolytic peptide was quantified for each protein using an internal standard (QconCAT). Data represent mean values over three biological repeats, each analyzed in three technical replicates. (C) Concentration of FtsA, MinD and MinE (most C-terminal peptide) as a function of the expression time course (from 0 to 6 hours). Symbols represent data from three independent experiments. The solid lines are fits to a mathematical model for gene expression kinetics (Methods section).

### Integration of MinDEC dynamic patterns and FtsA-FtsZ cytoskeletal structures on SLBs

We next investigated how the MinDEC and FtsA-FtsZ networks mutually interact on SLBs. The tricistron DNA was expressed in test-tube reactions to produce FtsA and MinDE. Purified FtsZ-A647 (FtsZ conjugated to the AlexaFluor647 fluorophore, 3 μM) and eGFP-MinC (fusion between the enhanced green fluorescent protein and MinC, 0.5 μM) were subsequently added, along with adenosine triphosphate (ATP, extra 2.5 mM) and guanosine triphosphate (GTP, extra 2 mM) (Fig. 2 A). Sample was transferred onto an SLB and the membrane organization of the FtsZ- and Min-subsystems was imaged by time-lapse fluorescence microscopy. We found that the membrane area close to the center of the chamber exhibited exclusively FtsA-FtsZ filaments and ring-like structures (Fig. 2 B), with cytoskeletal properties similar to that observed in the absence of Min proteins^20,30^. In contrast, the edge of the chamber was occupied by MinDEC dynamic patterns only. In between these two regions, mixed patterns involving the two subsystems were observed (Fig. 2 B). Spatial discrimination in distinct domains occurred immediately after supplying the PURE sample. We assume that molecular diffusion at the membrane is constrained by the edge of the chamber, resulting in a higher concentration of membrane-bound MinD and MinE^36,37^ that outcompete FtsA for membrane coverage.

**Figure 2.**
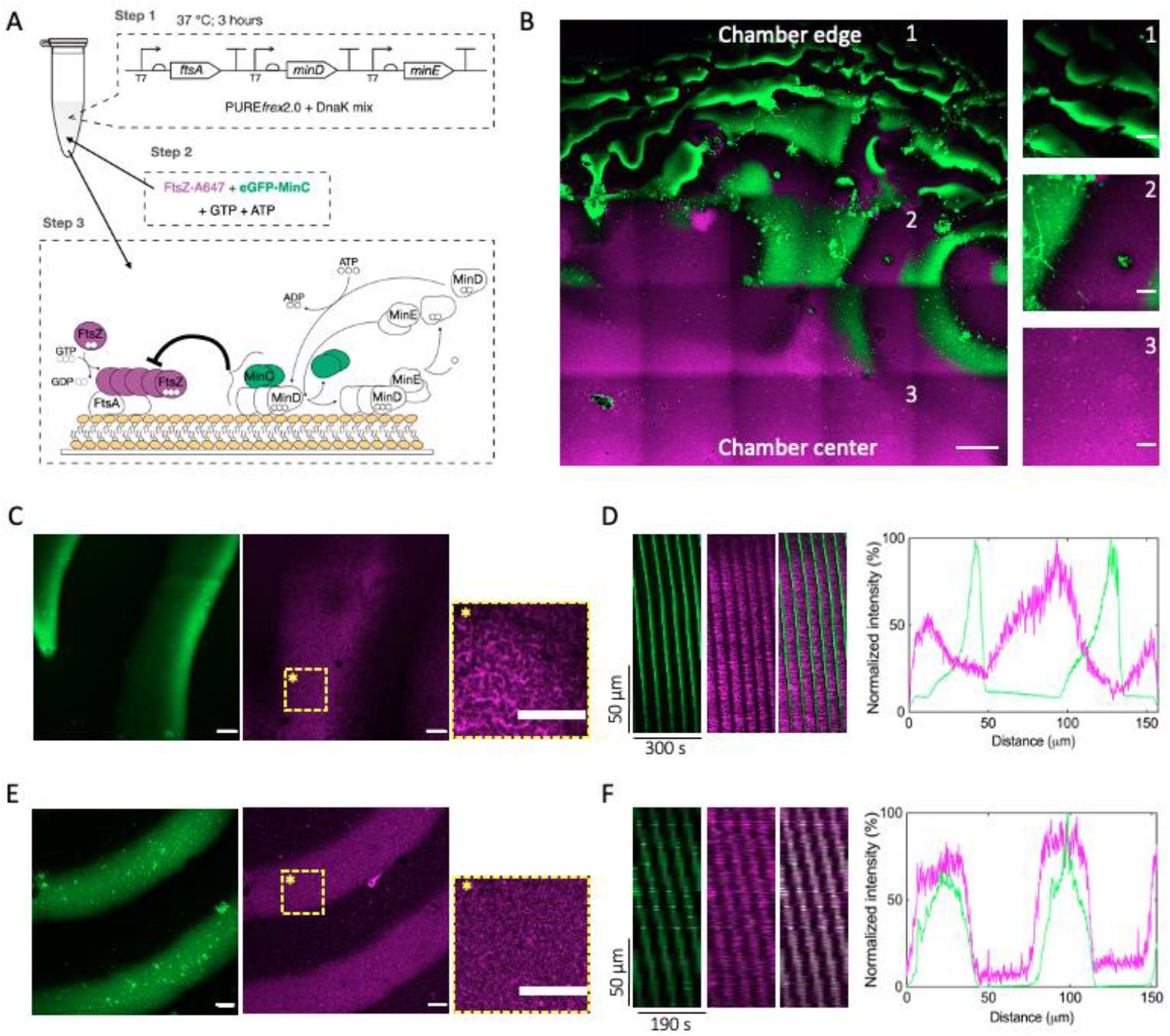
Integration of MinDEC surface waves with FtsA-FtsZ filament structures on supported membranes. (A) Schematic of the experimental approach. Genes *ftsA, minD* and *minE* were co-expressed from the three-cistron DNA template (5 nM) in PURE system with a DnaK chaperone mix for 3 hours at 37 °C. The solution was then supplemented with 2.5 mM ATP, 2.0 mM GTP, 0.5 μM eGFP-MinC and 3 μM FtsZ-A647 before transfer on top of an SLB. The cartoon depicts the main biochemical steps underlying FtsZ polymerization, its recruitment to the membrane by FtsA, and the influence of the reconstituted Min network. (B) Mosaic of 5×5 tile scan microscope images showing large-scale organization of FtsA-FtsZ and MinDEC dynamic patterns. FtsA-FtsZ rings mostly populate the center of the chamber (3), while MinDEC oscillating gradients dominate at the edges of the chamber (1). In between these two areas (2), correlated patterns of the two subsystems can be seen. Signals from eGFP-MinC and FtsZ-A647 are in green and magenta, respectively. Composite images of overlaid channels are shown. Scale bar is 50 μm in the mosaic image and 10 μm in the three zoomed-in images. (C) Fluorescence microscopy images of FtsA-FtsZ and MinDEC dynamic patterns taken from the intermediate SLB area (2). A zoom-in image of the framed area in the FtsZ channel is also displayed, showing the organization of FtsA-FtsZ into curved filaments and ring-like structures. Scale bars are 10 μm. (D) The time evolution of planar waves was analyzed and kymographs were constructed. On the right, examples of intensity profiles of MinC and FtsZ are shown. Color coding is the same as for the microscopy images. (E) Fluorescence microscopy images of FtsZ and MinDEC dynamic patterns when omitting FtsA. A zoom-in image of the framed area in the FtsZ channel is also displayed, showing that in the absence of FtsA, FtsZ fails to organize into cytoskeletal structures. (F) Kymographs constructed from the time series images shown in (E). On the right, intensity profiles of MinC and FtsZ signals showing that FtsZ travels with the MinDEC waves in the absence of a membrane anchor.

In the areas where FtsA-FtsZ and Min proteins coexisted, FtsZ could undergo different types of dynamic behaviors shaped by Min oscillatory gradients, including planar waves, rotating spirals, and standing waves (Fig. 2 B and C and Fig. S2). Fluorescence intensity profiles showed that membrane localization of MinC and FtsZ anti-correlated (Fig. 2D, Movie S1). The intensity distribution found for the traveling wave was broader for FtsZ than MinC. Calculated wavelength of MinC planar waves in the region where the two subsystems coexisted was 113 ± 30 μm and the wave velocity was 0.71 ± 0.30 μm s^−1^ (mean ± SD, from three biological replicates). Standing waves showed a characteristic oscillation time of 105 ± 24 s. In the area where FtsZ signal was not detectable, the Min wavelength was 57 ± 13 μm, i.e. about half the value measured in the intermediate region. This result suggests that membrane-bound FtsA-FtsZ filaments influence the Min wave properties by increasing the wavelength (Fig. S2 B). When omitting FtsA through expression of a bi-cistronic DNA template containing only the *minD* and *minE* genes, FtsZ rings were no longer observed on the bilayer (Fig. 2 E, Movie S2). Instead, FtsZ traveled in phase with MinC on the MinDE waves (Fig. 2 E and F, Movie S2). The observed colocalization of the MinC and FtsZ patterns was consistent with previous results that indicated weak transient interaction between MinC and non-membrane bound FtsZ^38^. Calculated wavelength was 59 ± 14 μm, similar to the value measured for MinDEC patterns in the area where FtsA-FtsZ were excluded. This result supports the hypothesis that the presence of FtsA-FtsZ on the membrane increases the wavelength of propagating Min waves.

Following a traditional view of the interplay between the two subsystems, the dynamic behavior of FtsZ reported in Fig. 2 B–C is expected to be the result of the inhibitory interaction of traveling MinC on FtsA-bound FtsZ. However, a different interpretation would be that FtsA-FtsZ oscillatory gradients originated from the direct action of the MinDE proteins, a possible scenario that we investigated in the next section.

### MinDE oscillatory gradients, without MinC, drive the formation of FtsA-FtsZ cytoskeletal patterns on SLBs

To explore the hypothesis that MinC has a dispensable function in regulating FtsA-FtsZ membrane dynamics, we repeated the experiments described above, this time replacing purified eGFP-MinC by a trace amount of purified eGFP-MinD (100 nM) as a reporter of the Min oscillations (Fig. 3 A). On a large scale, FtsA-FtsZ structures were more prominent in the center of the chamber, while Min waves preferentially populate the outer area of the chamber. Coexistence of both MinDE and FtsA-FtsZ dynamic patterns occurred in between these two regions (Fig. 3 B), similarly as observed in the presence of MinC (Fig. 2 B).

**Figure 3.**
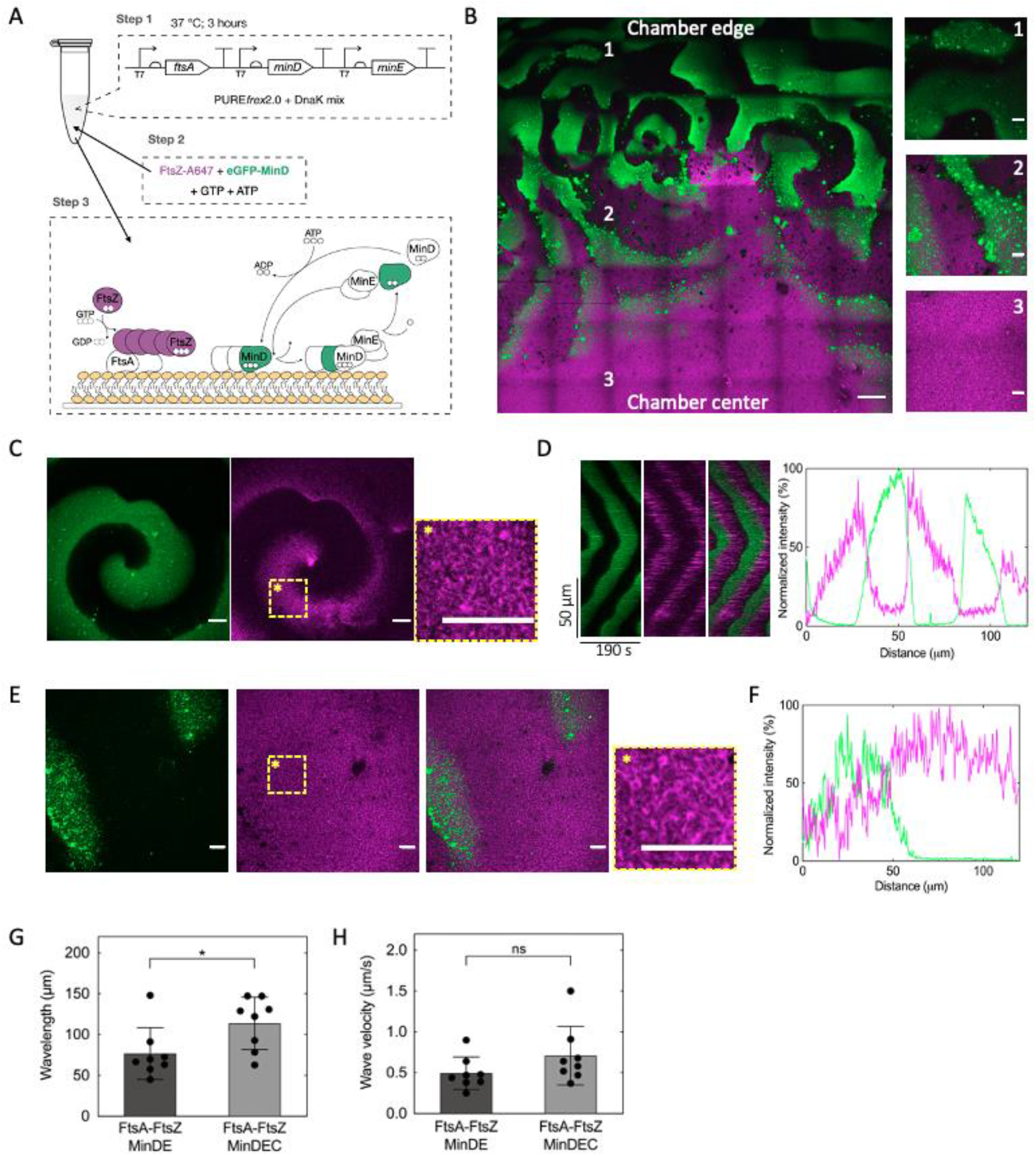
MinDE proteins, without MinC, regulate FtsA-FtsZ patterns on supported membrane assays. (A) Schematic of the experimental workflow for end-point expression assays. Genes *ftsA, minD* and *minE* were co-expressed from a single DNA template (5 nM) in the PURE system in the presence of DnaK chaperone mix for 3 hours at 37 °C. The solution was then supplemented with 2.5 mM ATP, 2.0 mM GTP, 100 nM eGFP-MinD and 3 μM FtsZ-A647 before transfer on top of an SLB. (B) Mosaic of 7×7 tile scan microscope images showing large-scale organization of FtsA-FtsZ and MinDE dynamic patterns. FtsA-FtsZ rings mostly populate the center of the chamber (3), while MinDE oscillating gradients dominate at the edges of the chamber (1). In between these two areas (2), correlated patterns of the two subsystems can be seen. Signals from eGFP-MinD and FtsZ-A647 are in green and magenta, respectively. Composite images of overlaid channels are shown. Scale bar is 50 μm in the mosaic image and 10 μm in the three zoomed-in images. (C) Fluorescence microscopy images of FtsA-FtsZ and MinDE dynamic patterns taken from the intermediate SLB area (2) showing sharp propagating waves of FtsA-FtsZ and MinDE on the SLB. A zoom-in image of the framed area in the FtsZ channel is also displayed, showing the organization of FtsA-FtsZ into curved filaments and ring-like structures. (D) The time evolution of the spiral wave in C was analyzed, and kymographs were constructed. On the right, examples of intensity profiles of MinD and FtsZ are shown. Color coding is the same as for microscopy images. (E) As in C but low-amplitude propagating waves of FtsZ anticorrelating with MinDE patterns are shown. (F) The time evolution of the wave was analyzed, and kymographs were constructed. On the right, examples of intensity profiles of MinD and FtsZ are shown. Color coding is the same as for microscopy images. (G, H) Graphs reporting the calculated wavelength (G) and velocity (H) for MinDEC and MinDE waves, both in the presence of FtsA-FtsZ. Data are from three biological replicates and two to three fields of view have been analyzed per sample. Bar height represents the mean value, and the standard deviation is appended. Symbols are values for individual fields of view aggregated from three biological replicates. Values obtained for different conditions were statistically compared by performing a two-tailed Welch’s *t*-test. Asterisk indicates *P* value <0.05, while ‘ns’ denotes a non-significant difference with *P* value >0.05.

Several dynamic behaviors were found in the extended area in which FtsA-FtsZ ring-like structures and MinDE coexisted. In most fields of view MinDE patterns could effectively rearrange FtsA-FtsZ filaments (Fig. 3 C and D and Fig. S3 A, Movie S3 and S4). In a few cases, anticorrelated propagating waves of FtsZ and MinDE with a low amplitude were also observed (Fig. 3E and F, Movie S4). These two types of rearrangements might be the manifestation of different local concentrations of MinDE or FtsA-FtsZ on the bilayer. Also, areas with a higher concentration of FtsA and FtsZ may support formation of more stable cytoskeletal structures (longer residence time on the membrane) that are less susceptible to be redistributed by MinDE. The fact that we did not observe such dim propagating waves of FtsZ in the presence of MinC (Fig. 2 B and C) can be explained by MinC’s depolymerizing effect that prevents stable FtsZ bundles from forming, reduces the local concentration of FtsA-FtsZ and facilitates propagation of the MinDE diffusion barrier.

Quantifying the Min wave properties in the areas where FtsA-FtsZ ring-like structures were effectively rearranged, we calculated a wavelength for the MinD signal of 76 ± 30 μm and wave velocity of 0.5 ± 0.2 μm s^−1^ (mean ± SD, from three biological replicates), while standing waves had a characteristic oscillation time of 109 ± 36 s. In regions with no visible FtsZ signal, a Min wavelength of 71 ± 22 μm was measured, with no significant differences between the center and the edge of the chamber (Fig. S3 B). In our previous work^24^, MinDE surface waves in PURE system exhibited a wavelength of 43 ± 7 μm and a velocity of 0.5 ± 0.1 μm s^−1^. The data reported here show an increased wavelength of the MinDE oscillations, suggesting that FtsA-FtsZ cytoskeletal structures influence the wave properties (Fig. S3 B). Another possibility could be the difference in MinDE concentrations in the two studies (4.2 ± 0.9 μM and 3.2 ± 0.4 μM in the present study vs 19 ± 7 μM and 5 ± 4 μM in the previous study, for MinD and MinE respectively). Moreover, we found that the wavelength increased in the presence of MinC, while the velocity remained mostly unaffected (Fig. 3 G and H).

We then asked whether FtsA alone, i.e. not engaged into FtsA-FtsZ co-filaments, was also subjected to spatial reorganization by the MinDE propagating diffusion barrier. Low-amplitude anticorrelated patterning of FtsA has already been reported with purified proteins in simple buffer conditions^26^. Here, a PURE solution containing pre-expressed FtsA, MinD and MinE was supplemented with purified FtsA conjugated to the AlexaFluor488 fluorophore (0.4 μM) for imaging (Fig. S3 C). MinDE waves were able to generate sharp FtsA dynamic patterns even in the absence of FtsZ. This finding corroborates previous observations with peripheral membrane proteins, but the modulation is noticeably more pronounced in our assay than previously reported^26^.

We performed a series of control experiments that confirm that MinDE constitutes the minimal set of proteins to re-organize FtsA-FtsZ cytoskeletal structures on SLBs (Fig. S3 D and E). We excluded the possibility that contaminating amounts of MinC originating from purification carry over in the eGFP-MinD sample could influence the results by performing the same experiment as in Fig. 3 B, this time omitting eGFP-MinD and using FtsZ-A647 as the only fluorescent reporter. Similar FtsZ oscillatory gradients were observed (Fig. S3 D), confirming our finding that MinDE constitutes the minimal set of proteins to re-organize FtsA-FtsZ cytoskeletal structures on SLBs. In another control, FtsA was omitted by expressing a DNA template encoding only for MinD and MinE. FtsZ was not recruited to the membrane, neither within nor outside the MinDE patterns (Fig. S3 E). The measured wavelength was 46 ± 11 μm (Fig. S3 B), further indicating that the presence of FtsA-FtsZ on the membrane increases the Min wavelength.

In addition, we demonstrated that in situ protein biogenesis through expression of the three-gene construct on top of the SLB supports the formation of MinDE(C) traveling waves that pattern FtsA-FtsZ co-filaments (Fig. S4). The two subsystems coexist over the entire membrane surface, which contrasts with the spatial segregation of Min waves and FtsA-FtsZ filaments when pre-expressed proteins were added to the imaging chamber (Fig. 2 B and Fig.3 B).

### MinC is dispensable for coupling the Min waves and FtsA-FtsZ in closed compartments

We finally investigated if the findings from flat bilayer assays were also valid in a closed environment that better mimics the cellular context in terms of membrane surface-to-volume ratio and finite protein numbers. To do so, the FtsA-FtsZ and Min systems were co-reconstituted within lipid microdroplets. We first expressed the tri-cistronic DNA template in a test tube and the pre-ran PURE*frex*2.0 solution was encapsulated inside water-in-oil droplets (Fig. 4 A). The vast majority of the droplets displayed both FtsZ and Min protein signals on the inner surface (Fig. S5A and S6A, Movie S5). Only a few droplets exhibited either Min dynamic patterns or FtsA-FtsZ networks. Time lapse imaging showed that in both samples with (Fig. 4 B, Fig. S5 and Movie S6) and without MinC (Fig. 4 C, Fig. S6 and Movie S6), dynamic concentration gradients of Min and FtsZ formed at the lipid monolayer, and that the two patterns were anticorrelated, as observed on SLBs. In most of the droplets, FtsZ and MinDE(C) assembled on opposite sides of the membrane forming two or multiple domains (Fig. 4 B and C, Figs. S5 A,C and S6 A,C). These patches were not static, they migrated along the droplets surface (Fig. 4 B and C, Movie S6). Other droplets exhibited circling patterns in which MinDE(C) and FtsZ chased one another along the droplet interface, with no visible intraluminal diffusion within the imaging plane (Fig. 4 B and Movie S6). In droplets larger than 20 μm in diameter, it was possible to visualize cytoskeletal FtsZ structures forming a cortex and being re-localized by Min waves (Fig. 4 D and Fig. S5 D and S6 D and Movie S7). More complex dynamic patterns were also observed in large droplets, including multiple interfacial FtsZ polarization sites rearranged by the Min dynamics (Fig. 4 E and Fig. S5 and S6), resembling the standing Min waves reported on SLBs. For the different phenotypes the antiphase localization of the two subsystems persisted in time. Parameter values for the pattern dynamics were broad, with wavelength and wave velocity respectively ranging between 42-174 μm and 0.16-0.58 μm s^−1^ (8 droplets) with MinC, and between 24-167 μm and 0.23-0.60 μm s^−1^ (8 droplets) without MinC. These values were consistent with those found on SLB. No significant differences due to the presence or absence of MinC could be measured.

**Figure 4.**
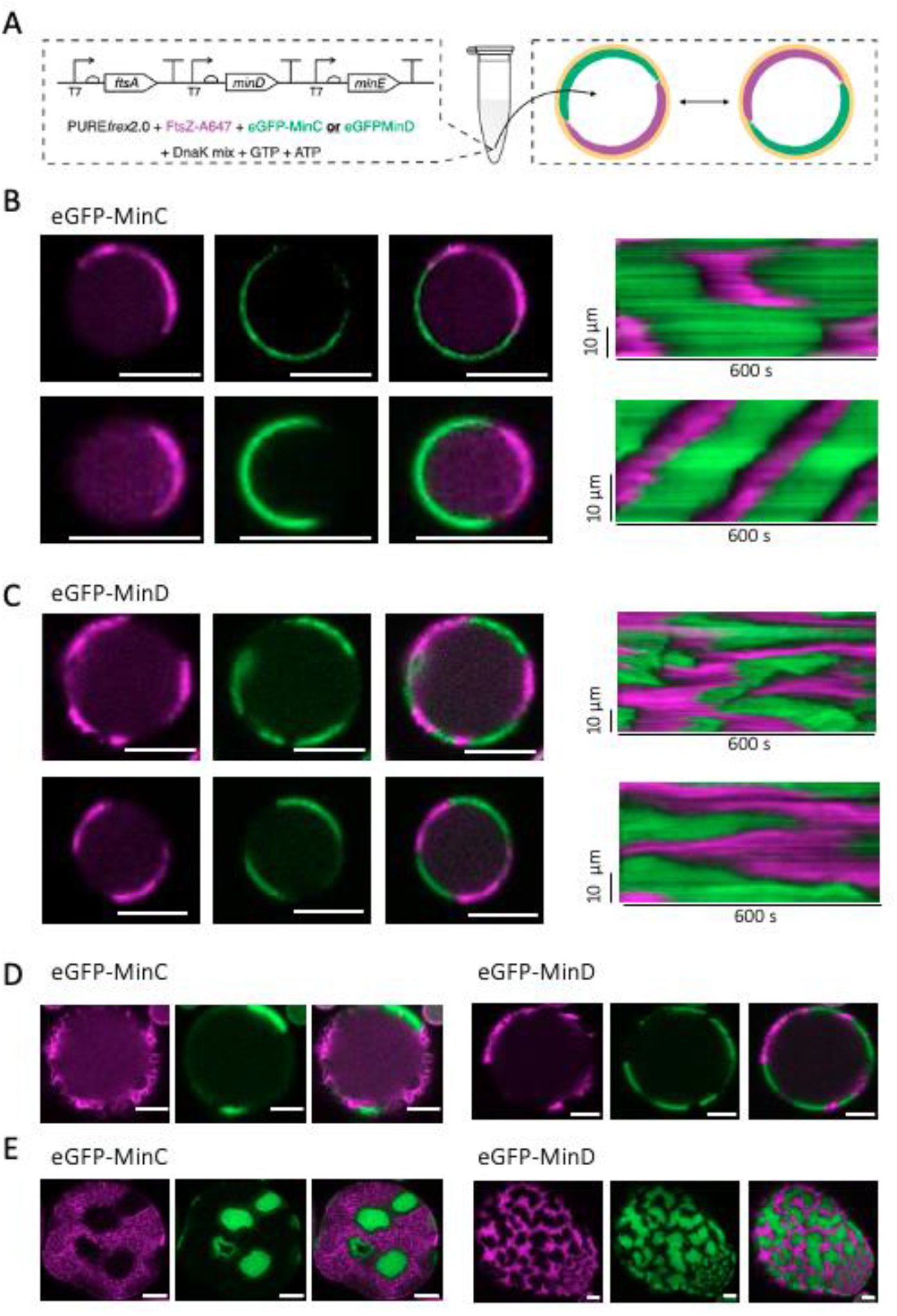
Functional reconstitution of MinDE(C) and FtsA-FtsZ networks in microdroplets. (A) Schematic illustration of the droplet assays. The DNA template containing the genes *ftsA*, *minD*, and *minE* (5 nM) was expressed in a test tube in the presence of DnaK mix. The pre-ran PURE*frex*2.0 solution was encapsulated inside water-in-oil droplets, along with ATP (2.5 mM), GTP (2 mM), FtsZ-A647 (3 μM) and either purified eGFP-MinC (0.5 μM) or eGFP-MinD (100 nM). (B) Fluorescence images (split channels and composite) of two droplets exhibiting antiphase dynamic patterns of MinDEC and FtsA-FtsZ. The corresponding kymographs are displayed. The droplet at the bottom exhibited a circling pattern yielding a characteristic kymograph. (C) Same as in B, except that eGFP-MinC was substituted with eGFP-MinD. (D) Same as in B and C (exact condition is as specified) except that bigger droplets (diameter >20 μm) were imaged. Here, more defined cytoskeletal FtsZ structures were visible. (E) Same as in B-D (exact condition is as specified) except that the images have been acquired closer to the dome of large droplets. Here, multiple interfacial FtsZ polarization sites rearranged by the Min dynamics were visible. Color coding: eGFP-MinC and eGFP-MinD (green), FtsZ-A647 (magenta). Scale bars are 10 μm.

These data confirm the functional integration of FtsA-FtsZ and Min proteins in closed compartment, further demonstrating that MinC is not essential to create dynamic FtsZ patterns in vitro when FtsA acts as the membrane anchor.

## Discussion

In this work we reconstituted MinDE(C) dynamics with FtsA-FtsZ cytoskeletal networks in a cell-free environment that accounts for the complex molecular content of the bacterial cytoplasm. To accomplish this, we used the PURE system to express a three-gene DNA template encoding for MinD, MinE and FtsA (Fig. 1). FtsA is an essential, widely conserved membrane anchor protein, playing a major role in Z-ring formation and downstream protein recruitment^7^. It is known that the nature of the FtsZ membrane anchor influences the dynamics of FtsZ assembly in SLB assays^20^. Therefore, the questions of whether and how the Min system drives the spatial rearrangement of FtsA-bound FtsZ are relevant; yet, they have hitherto not been explored.

We found that membrane-bound FtsA-FtsZ filaments organize into dynamic, anti-phased, patterns driven by the MinDEC system (Fig. 2 B and C and Fig. S4 A). The traditional explanation for the emergence of FtsZ oscillation gradients is that the membrane-tethered polymers reorganize as a result of the interaction with MinC(D). However, similar dynamic behaviors were observed in the absence of MinC (Fig. 3 B and C and Fig. S4 B and C), demonstrating that MinDE act as a minimal spatiotemporal regulator of FtsA-anchored FtsZ filaments on SLBs. This conclusion is also valid when the two subsystems are enclosed in water-in-oil droplets (Fig. 4).

FtsZ redistribution on lipid membranes has not been documented in the absence of MinC when ZipA (soluble version) was used in place of FtsA^38^. Dynamic patterns of ZipA-FtsZ only emerged with MinC traveling on MinDE waves. In this previous study, ZipA was not mobile^38^, which might have compromised the rearrangement of ZipA-FtsZ cofilaments by the MinDE oscillatory gradients. When the Min system was combined with the fusion protein FtsZ-YFP-MTS on SLBs, two distinct scenarios were reported^26^: dynamic FtsZ-YFP-MTS rings were reorganized by MinDE, while static FtsZ-YFP-MTS networks remained unaffected. In both cases, supplementing MinC stimulated formation of clear FtsZ-YFP-MTS patterns^26^. Moreover, FtsZ-YFP-MTS and Min systems have been coupled inside droplets^39^ or PDMS compartments^40^, showing that Min waves comprising MinC create dynamic FtsZ surface patterns^39,40^. These earlier results involving ZipA^38^ or FtsZ-YFP-MTS^26,39,40^ differ from the robust dynamic patterning of FtsA-FtsZ reported here in the absence of MinC, indicating that different membrane targeting proteins may respond differently to Min concentration gradients. Based on the present observations, both on open and closed membranes, we propose that MinDE oscillations alone constitute the minimal localization mechanism of the FtsA-FtsZ proto-ring. This assumption is reinforced by the recent discovery that MinDE act as a general propagating diffusion barrier exerting steric pressure that can redistribute membrane proteins in vitro^26,27^.

In vivo, functional MinC is undoubtedly needed to prevent septum misplacement and asymmetric division, a phenotype known as minicell^10^. The dispensable function of MinC in regulating FtsZ-FtsA dynamics, as reported here, suggests that MinDE oscillations play a more central role in patterning critical division components in *E. coli* than solely distributing MinC at the cell pole, where Z-ring formation is prohibited. After all, in vivo studies have shown that ZipA, as well as other early division proteins, counter-oscillate with MinC^41^. ZipA traveling in phase with FtsZ was interpreted as resulting from the interaction between these two proteins during filament depolymerization by MinC. However, ZipA might also be redistributed by the MinDE oscillations. FtsA and ZipA follow the FtsZ pattern in time, and the three proteins are detectable at midcell with no significant delay^31^, suggesting a tightly coordinated regulatory mechanism. The earliest phase of cytokinesis in *E. coli* includes the cytosolic assembly of short FtsZ protofilaments that bind transiently everywhere on the inner membrane and, over time, migrate and cluster at the cell’s center^42^. Our results suggest that MinDE promotes the localization of short FtsA-FtsZ filaments at midcell by acting as a propagating steric hindrance on the membrane, competing with the fast detaching-binding short filaments (Fig. 5 A). The high local concentration of short filaments in the middle of the cell then encourages lateral interaction, bundling and extension of filaments, and eventually proto-ring formation. It should be noted than in slow growing *E. coli* cells, condensation of membrane-bound FtsZ filaments into a ring occurs in the absence of the Min system^43^, indicating that other mechanisms guide the ring assembly and its localization. As shown in Fig. 3 E, in some instances FtsA-FtsZ structures cannot effectively be displaced by MinDE only. MinC inhibits long FtsA-FtsZ filaments by depolymerizing membrane-bound FtsZ structures, by competing with the membrane proteins for binding to FtsZ^44^, and by capping growing polymers^45^. By lowering the residence time of FtsA-FtsZ filaments on the membrane, MinC may assist MinDE in effectively reorganizing the FtsA-FtsZ structures (Fig. 5 B). In addition, MinC could be critical in managing the pool of FtsZ that is not at the division site (up to 70% of the available FtsZ^46^), i.e. by buffering the concentration of cytosolic FtsZ. The observation that FtsZ travels in phase with MinC in the absence of a membrane anchor lends support to this hypothesis (Fig. 2 E). In *E. coli*, MinC overexpression impairs cytokinesis, resulting in cell filamentation^47^. The inhibitory effect is caused by MinC sequestering a large pool of free FtsZ^48^. This depletion process is likely enhanced by the interaction of MinC to MinD, which increases the local concentration of MinC at the membrane.

**Figure 5.**
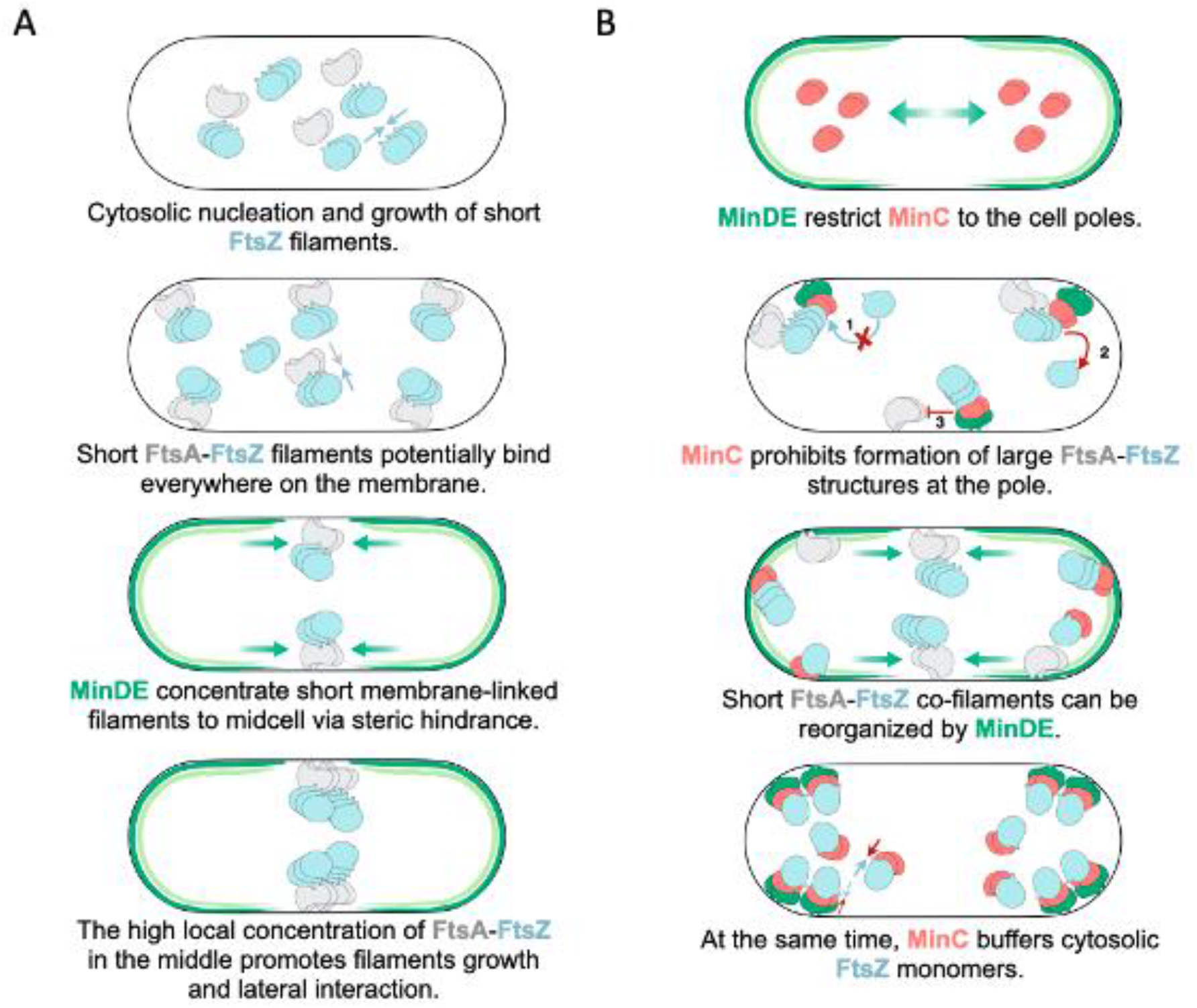
Model for the patterning of bacterial cytokinesis components by MinDE oscillations in *E. coli*. (A) Short protofilaments of FtsA-FtsZ nucleate and grow in the cytosol. The short FtsZ-FtsA (and possibly ZipA-FtsZ) filaments bind at random locations along the membrane. MinDE propagating diffusion barrier concentrates the short filaments at midcell. The high local concentration of short filaments in the middle promotes lateral interaction, bundling and extension of filaments, and subsequent proto-ring formation. (B) MinD-MinE dynamics generate a concentration maximum of MinC at the poles, where the formation of extended FtsA-FtsZ structures that could not be displaced by MinDE only is prohibited by the following processes: MinCD inhibits long FtsA-FtsZ structures by capping growing FtsZ polymers (1), by depolymerizing FtsZ structures (2), and by competing with other membrane proteins for binding to FtsZ (3). The short co-filaments have a reduced lifetime at the membrane and can more easily be outcompeted by MinDE at the division site. Furthermore, MinC is responsible for sequestering FtsZ monomers through direct interaction, keeping the cytosolic concentration of FtsZ lower than the threshold to form long filaments and bundles. The interaction between membrane-bound MinD and MinC boosts the local concentration of MinC, which might operate as a mechanism to accentuate the depletion of FtsZ subunits. Together, the processes described in A and B restrict proto-ring formation to midcell.

Finally, our finding will have implications toward the reconstitution of a more elementary regulatory mechanism for positioning the division site in a synthetic cell. DNA-encoded protein synthesis and compartmentalization are important design elements to building a minimal cell. Therefore, demonstrating that the FtsZ and Min subsystems can be partly expressed from a multigene DNA template, functionally integrated, and encapsulated in droplets is a step forward to establishing a synthetic division mechanism. Implementation of a coupled FtsA-FtsZ-MinDE(C) system inside deformable lipid vesicles^24,30^ might be a route to symmetrical division of artificial cells. Spatiotemporal redistribution of other bacterial division proteins, such as ZapA, ZapB, MatP, and of lipid synthesis enzymes49 could also be explored in the future.

## Materials and Methods

### Purified proteins

The proteins eGFP-MinD and eGFP-MinC were purified using previously described methods ^23^. Protein concentrations were quantified via Bradford assay and eGFP absorbance measurements. FtsZ was dialyzed against 20 mM Hepes/HCl, pH 8.0, with 50 mM KCl, 5 mM MgCl_2_, and 1 mM ethylenediaminetetraacetic acid (EDTA). The protein was initially polymerized at 30 °C using 20 mM CaCl_2_ and 2 mM GTP, then the mixture was incubated at 30 °C for 15 min with a 20-fold excess of Alexa Fluor 647 (A647). This two-step process helps reducing labeling-induced alterations in FtsZ assembly. The precipitate was resuspended on ice in 50 mM Tris/HCl, pH 7.4, with 100 mM KCl, and the unbound fluorescent probe was extracted by gel filtration, as per standard protocol for column purification (Gel Wizard SV). Purified FtsZ-Alexa647 was kept in storage buffer (50 mM Tris, 500 mM KCl, 5 mM MgCl_2_, and 5% glycerol) at pH 7, as previously described^50^. FtsA was expressed and purified according to published protocols^51^. Cells were suspended in 50 mM Tris-HCl, 1 mM TCEP, 1 mM PMSF, pH 7.5, treated with 10 g/mL DNase, sonicated and centrifuged. The pellet was washed two times with 20 mM Tris-HCl, 10 mM EDTA, pH 7.5, and 1% (v/v) Triton X-100. Inclusion bodies were dissolved in 20 mM Tris-HCl, 5 M guanidine-HCl, 0.5 M NaCl, pH 7.5, loaded in a HisTrap FF column (GE Healthcare) and rinsed in 20 mM imidazole. An imidazole elution gradient (20–500 mM) was applied and the eluted fractions were kept at −80 °C. FtsA was refolded by dialysis against 50 mM Tris-HCl, 0.5 M NaCl, 5 mM MgCl_2_, 0.2 mM TCEP, and 0.1 mM ADP, pH 8 at 4 °C. Buffer was replaced with 50 mM Tris-HCl, 0.5 M KCl, 5 mM MgCl_2_, 0.2 mM TCEP and 0.1 mM ADP, pH 7.5. Concentration of FtsA was measured by Bradford assay. FtsA was fluorescently labeled with AlexaFluor-488 (1:10 molar ratio) for 30 min in 50 mM Hepes/HCl (pH 8.0), 100 mM KCl, and 5 mM MgCl_2_.

### Preparation of DNA constructs

*minD,minE* and *FtsA* genes were amplified by standard polymerase chain reaction (PCR) amplified using Phusion High-Fidelity DNA polymerase (New England Biolabs, USA) as previously reported^24,30^. The plasmid DE was assembled via Gibson assembly (Gibson Assembly Master Mix of New England BioLabs, Inc.). The assembly was performed at equimolar concentrations of linearized plasmid MinDpUC57 and MinE DNA fragments for 1 hour at 50 °C. The primers in Table S1 were used: ChD173+504 and ChD1139+1140. Transformation of the Gibson assembly products into *E. coli* TOP10 competent cells was done by heat shock, after which cells were resuspended in 200 μL of fresh prechilled liquid lysogeny broth (LB) medium and incubated for 1 hour at 37 °C and 250 r.p.m. Then, the cultures were plated in solid LB medium with ampicillin and grew overnight at 37 °C. Colonies were picked up and cultured in 1 mL of liquid LB medium with 100 μg mL^−1^ of ampicillin for 16 hours at 37 °C and 250 r.p.m. Plasmid purification was performed using the PureYieldTM Plasmid Miniprep System (Promega). Concentration and purity of DNA were checked on NanoDrop. Linear DNA construct was prepared by PCR using the forward and reverse primers ChD365 and ChD174, respectively, annealing to the T7 promoter and T7 terminator sequences (Supplementary Table1).

The plasmid ADE was assembled via Gibson assembly, using equimolar concentrations of linearized plasmid FtsApU57 and the MinDE DNA fragment, for 1 hour at 50 °C. The primers in Table S1 were used: ChD420+1145 and ChD365+174. Transformation of the Gibson assembly products into *E. coli* TOP10 competent cells was done by heat shock, after which cells were resuspended in 200 μL of fresh prechilled liquid lysogeny broth (LB) medium and incubated for 1 hour at 37 °C and 250 r.p.m. Then, the cultures were plated in solid LB medium with ampicillin and grew overnight at 37 °C. Colonies were picked up and cultured in 1 mL of liquid LB medium with 100 μg mL^−1^ of ampicillin for 16 hours at 37 °C and 250 r.p.m. Plasmid purification was performed using the PureYieldTM Plasmid Miniprep System (Promega). Concentration and purity of DNA were checked on NanoDrop. Linear DNA construct was prepared by PCR using the forward and reverse primers ChD1187 and ChD174, respectively, annealing to the T7 promoter and T7 terminator sequences. For both linear constructs (DE and ADE) amplification products were checked on a 1% agarose gel and were further purified using Wizard SV gel (standard column protocol). Concentration and purity were measured by NanoDrop. All sequences were confirmed by sequencing. MinE-, and FtsA-coding DNA fragments (starting with a T7 promoter and ending with a T7 terminator) are sequence-optimized for codon usage, CG content, and 5ʹ mRNA secondary structures. Sequences of the linearized constructs can be found in the Supplementary Methods.

### Cell-free gene expression in bulk

Gene expression was performed using PURE*frex*2.0 (GeneFrontier Corporation, Japan) and 5 nM of a linear DNA template following the supplier’s recommendations. The solution was supplemented with 1 μL of DnaK Mix (GeneFrontier Corporation), which consists of purified DnaK, DnaJ, and GrpE chaperone proteins from *E. coli*. Twenty microliter reactions were run in PCR tubes for 3 hours at 37 °C. Purified proteins (FtsZ-A647, eGFP-MinC/D), ATP and/or GTP were supplied to the mixture only when specified.

### QconCAT purification

A quantitative concatemer (QconCAT) protein was designed to contain two specific peptides for FtsA and MinD, one for MinE and two for ribosomal core proteins. These include the most C-terminal proteolytic peptide that we could accurately identify for each protein of interest^24,30^. QconCAT was expressed in BL21(DE3) cells in M9 medium with ^15^NH_4_Cl and ampicillin. A pre-culture was diluted 1:100 to a 50 mL expression culture. Protein expression was induced at OD_600_ = 0.5 with 1 mM isopropyl β-d-1-thiogalactopyranoside and cells were grown for 3 hours at 37 °C. Cells were harvested by centrifugation and the pellet was dissolved in 1 mL B-PER. 10 μL of 10 mg mL^−1^ lysozyme and 10 μL of DNaseI (ThermoScientific, 1 U μL^−1^) were added and the sample was incubated for 10 min at room temperature. The lysate was centrifuged for 20 min at 16,000 × *g* and the pellet resuspended in 2 mL of a 1:10 dilution of B-PER in MilliQ water. The sample was twice again centrifuged, and the pellet was resuspended in 2 mL 1:10 diluted B-PER and centrifuged again. The pellet was resuspended in 600 μL of 10 mM Tris-HCl pH 8.0, 6 M guanidinium chloride and incubated at room temperature for 30 min. After spinning down the insolubilized protein fraction the supernatant was loaded onto an equilibrated mini NiNTA spin column (Qiagen) and the flow-through was reloaded twice to maximize protein binding. The column was washed twice with 600 μL of 10 mM Tris-HCl pH 6.3, 8 M urea and the QconCAT was eluted with 3 × 200 μL of 10 mM Tris-HCl pH 4.5, 8 M urea, 400 mM imidazole. The eluate was dialyzed overnight and for additional 4 hours against 10 mm Tris-HCl pH 8.0, 100 mM KCl with a 10-kDa cutoff slide-a-lyzer casette (ThermoScientific).

### Trypsin digest

Per LC-MS injection, 1.5 μL of PURE system reaction was mixed with 3 μL of 100 mM Tris-HCl pH 8.0, 0.3 μL of 20 mM CaCl_2_, and 0.8 μL MilliQ water. Samples were incubated at 90 °C for 10 min to stop the reaction. Then, 0.6 μL of QconCAT (0.3 mg mL^−1^) was added, the sample was incubated again at 90 °C for 10 min and after cooling to room temperature, 1 μL 250 mM fresh iodoacetamide was added and the sample incubated for 30min at room temperature in the dark. Then 0.3 μL of 1 mg mL^−1^ trypsin (trypsin-ultra, MS-grade, New England Biolabs) was added and samples were incubated at 37 °C overnight. After addition of 0.7 μL 10% trifluoroacetic acid, samples were centrifuged in a table-top centrifuge (5415 R, Eppendorf) for 10 min at maximum speed. The supernatant was transferred to a glass vial with small-volume insert for LC-MS/MS analysis.

### LC-MS/MS analysis

LC-MS/MS analysis was performed on a 6460 Triple Quad LCMS system (Agilent Technologies, USA) using Skyline software^52^. Per run 7 μL of sample was injected to an ACQUITY UPLC® Peptide CSH™ C18 Column (Waters Corporation, USA). The peptides were separated in a gradient of buffer A (25 mM formic acid in MilliQ water) and buffer B (50 mM formic acid in acetonitrile) at a flow rate of 500 μL per minute and at a column temperature of 40 °C. The column was equilibrated with 98% buffer A. After injection, the gradient was changed linearly over 20 min to 70% buffer A, over the next 4 min to 60% buffer A, and over the next 30 s to 20% buffer A. This ratio was held for another 30 s and the column was finally flushed with 98% buffer A to equilibrate for the next run. Selected peptides were measured by multiple reaction monitoring. For both FtsA and MinD, two peptides were analyzed, one for MinE. Two peptides from ribosomal proteins were also measured as a control. The peak area ratio of unlabeled peptides (^14^N, expressed or PURE component) and ^15^N-labeled QconCAT peptides was calculated for the ribosomal core proteins (average value over the two peptides) and for the proteins of interest. The concentration of the selected peptides for the proteins of interest was then deduced knowing the concentration of ribosomal core proteins in PURE system. Data processing was performed using a script written in Mathematica (Wolfram Research, version 11.3).

### Phenomenological fitting of FtsA, MinD and MinE production kinetics

FtsA, MinD, and MinE most C-terminal peptide concentrations were fit using a phenomenological model to estimate apparent kinetic parameters: final yield, production rate, and translation lifetime. The following sigmoid equation was used^53^:

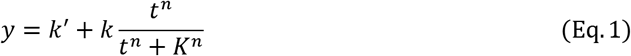

where *t* is the time in minutes, *y* the peptide concentration at a given time point, and *k*′, *k*, *K*, and *n* are fit parameters. Using this expression, the final yield corresponds to *k* and the plateau time, or expression lifespan, is expressed as:

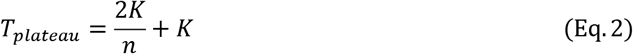

The apparent translation rate, which is defined as the steepness at time *t* = *K*, is:

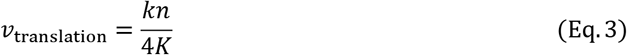

The kinetics from three independent experiments were fit, and the extracted parameter values are reported as the average and standard deviation.

### Fabrication and cleaning of the imaging chamber

Home-made imaging glass chambers were used in all the assays ^24^. Three microscopy glass slides were bonded together using NOA 61 glue (Norland Products). Holes were drilled across the three-slide layer, with diameters of 2.5 mm. A150-μm thick coverslip (Menzel-Gläser) was attached on one side of the slide with NOA 61 to form the bottom of the chamber. Chambers were cleaned by 10-min washing steps in a bath sonicator (Sonorex Digitec, Bandelin) using the following solutions: chloroform and methanol (volume 1:1), 2% Hellmanex, 1 M KOH, 100% ethanol, and MilliQ water. In addition, after a couple of experiments the glass chambers were subjected to acid Piranha treatment.

### Lipids

1,2-dioleoyl-sn-glycero-3-phosphocholine (DOPC), 1,2-dioleoyl-sn-glycero-3phosphoglycerol (DOPG) and 1′,3′-bis[1,2-dioleoyl-sn-glycero-3-phospho]-glycerol (18:1 CL) were purchased from Avanti Polar Lipids as chloroform solutions.

### Preparation of small unilamellar vesicles

Small unilamellar vesicles (SUVs) were used as precursors for SLB production. DOPC (4 μmol) and DOPG (1 μmol) lipids dissolved in chloroform were mixed in a glass vial. A lipid film was formed on the vial wall by solvent evaporation under a moderate flow of argon and desiccated for 30 min at room temperature. The lipid film was resuspended with 400 μL of SLB buffer (50 mM Tris, 300 mM KCl, 5 mM MgCl_2_, pH 7.5) and the solution was vortexed for a few minutes. The final lipid concentration was 1.25 mg mL^−1^. A two-step extrusion (each of eleven passages) was performed utilizing an Avanti mini extruder (Avanti Polar Lipids) with 250 μL Hamilton syringes, filters (drain disc 10 mm diameter, Whatman), and a polycarbonate membrane with pore size 0.2 μm (step 1) or 0.03 μm (step 2) (Nuclepore track-etched membrane, Whatman).

### Formation of SLBs

The imaging chamber was treated with oxygen plasma (Harrick Plasma basic plasma cleaner) for 30 min to activate the glass surface. Immediately after, an SUV solution was added to the sample reservoir at a final lipid concentration of 0.94 mg mL^−1^ together with 3 mM CaCl_2_. The chamber was sealed with a 20×20 mm coverslip by using a double-sided adhesive silicone sheet (Life Technologies). After sample incubation at 37 °C for 30 min, the chamber was carefully opened and the SLB was rinsed six times with SLB buffer.

### Activity assays on supported membranes

The constructs *ftsA*-*minD*-*minE* and *minD*-*minE* were expressed with PURE*frex*2.0 in test tubes as described above and the solution was supplemented with the following compounds: 2 mM GTP, 2.5 mM ATP, 3 μM purified FtsZ-A647, and either 0.5 μM eGFP-MinC or 100 nM purified eGFP-MinD as specified in the text (all final concentrations) in a total volume of 20 μL. The sample was added on top of an SLB, and the imaging chamber was sealed with a 20×20 mm coverslip by using a double-sided adhesive silicone sheet. For assays involving in situ expression, a 20 μL PURE*frex*2.0 mixture containing one of the two DNA constructs was supplemented with 2 mM GTP, 2.5 mM ATP, 3 μM purified FtsZ-A647, and either 0.5 μM eGFP-MinC or 100 nM purified eGFP-MinD as specified in the text (all final concentrations), and the solution was directly incubated on top of an SLB. 0.4 μM purified FtsA-A488 was used only when specified in the main text. The chamber was closed as indicated above, and the sample was immediately imaged with a time lapse fluorescence microscope for up to 5.5 hours at 37 °C. Numerous fields of view were acquired at various time points during the expression period.

### Droplet preparation

15 mol% 18:1 CL and 85 mol% DOPC in chloroform were mixed in a glass vial. A lipid film was deposited on the wall of the vial upon solvent evaporation through a gentle flow of nitrogen and was further desiccated for 30 min at room temperature. The lipid film was resuspended with mineral oil (Sigma Aldrich, St. Louis, MO) to a final concentration of 2.5 mg mL^−1^ and the solution was vortexed for a few minutes. The inner solution consisted of a pre-ran PURE system solution supplemented with DnaK, 2 mM GTP, 2.5 mM ATP, 3 μM purified FtsZ-A647, and either 0.5 μM eGFP-MinC or 100 nM purified eGFP-MinD as indicated in the text. Five microliter of inner solution was added to 30 μL of the oil/lipid mixture. The droplet-oil mixture was pipetted on a coverslip and imaged with a fluorescence confocal microscope.

### Confocal microscopy

Droplets and SLBs were imaged using a Nikon A1R Laser scanning confocal microscope with an SR Apo TIRF 100× oil-immersion objective. FtsZ-A647 and eGFP-MinC/D were excited using the 640-nm and 488-nm laser lines, respectively, and suitable emission filters were used. For image acquisition, the software NIS (Nikon) was utilized, with identical settings for all samples. Throughout imaging, samples were kept on a temperature-controlled stage (Tokai Hit INU) that was held at 37 °C.

### Image analysis

Fiji^54^ or MatLab version R2020b were used for image analysis. Fiji’s profile plots tool was used to generate fluorescence intensity profiles. The tool displays a two-dimensional graph of pixel intensities along a line drawn in the direction of the moving wave. The wavelength and velocity of the dynamic patterns on SLBs were determined by producing a kymograph parallel to the direction of the traveling wave. Lines were identified using a linear Hough transform after binarization of the kymograph using Sobel edge detection. The slope of the lines equates to the wave velocity, while the distance between the lines corresponds to the wavelength. The frequency of oscillations of a standing wave was determined by computing the autocorrelation function over time for each pixel and identifying the first peak of the function.

### Data availability

All materials, detailed protocols and the Mathematica script for LC-MS/MS data analysis are available on the request from the corresponding author.

## Supplementary materials

SI information and the extended methods sections contain sequences, primers, and the transitions of the MS/MS measurements for the proteolytic peptide analysis.

## Acknowledgments

We thank Ilja Westerlaken for purifying the eGFP-MinD and eGFP-MinC proteins and the group of Petra Schwille for providing the corresponding expression plasmids. We also thank Germán Rivas and Mercedes Jimenez for providing us with the purified FtsZ and FtsA and for discussing the data, and Jaan Mannik for fruitful discussions. Microscopy measurements were performed at the Kavli Nanolab Imaging Center Delft. This work was financially supported by the Netherlands Organization for Scientific Research (NWO/OCW) through the “BaSyC – Building a Synthetic Cell” Gravitation grant (024.003.019).

## Author Contributions

EG and CD designed experiments and wrote the paper. EG and AD performed experiments and analyzed data. All the authors discussed the results.

