## Supplementary Information for "Min waves without MinC can pattern FtsA-FtsZ filaments on model membranes"

**Supplemental information includes:**

Extended methods
Figures S1 to S6
Tables S1 to S2
Legends for Movies S1 to S7

**Other supplementary materials for this manuscript:**

Movies S1 to S7 are provided as separate files

### Extended methods

#### DNA sequence of the *minD-minE* construct

TAATACGACTCACTATAGGGGAATTGTGAGCGGATAACAATTCCCCTCTAGAAATAATTTTGTTTAACTT
TAAGAAGGAGATATACATATGGCACGCATTATTGTTGTTACTTCGGGCAAAGGGGGTGTTGGTAAGACA
ACCTCCAGCGCGGCCATCGCCACTGGTTTGGCCCAGAAGGGAAAGAAAACTGTCGTGATAGATTTTGA
TATCGGCCTGCGTAATCTCGACCTGATTATGGGTTGTGAACGCCGGGTCGTTTACGATTTCGTCAACGT
CATTCAGGGCGATGCAACGCTAAATCAGGCGTTAATTAAAGATAAGCGTACTGAAAATCTCTATATTCTG
CCGGCATCGCAAACACGCGATAAAGATGCCCTCACCCGTGAAGGGGTCGCCAAAGTTCTTGATGATCT
GAAAGCGATGGATTTTGAATTTATCGTTTGTGACTCCCCGGCAGGGATTGAAACCGGTGCGTTAATGGC
ACTCTATTTTGCAGACGAAGCCATTATTACCACCAACCCGGAAGTCTCCTCAGTACGCGACTCTGACCG
TATTTTAGGCATTCTGGCGTCGAAATCACGCCGCGCAGAAAATGGCGAAGAGCCTATTAAAGAGCACCT
GCTGTTAACGCGCTATAACCCAGGCCGCGTAAGCAGAGGTGACATGCTGAGCATGGAAGATGTGCTG
GAGATCCTGCGCATCAAACTCGTCGGCGTGATCCCAGAGGATCAATCAGTATTGCGCGCCTCTAACCA
GGGTGAACCGGTCATTCTCGACATTAACGCCGATGCGGGTAAAGCCTACGCAGATACCGTAGAACGTC
TGTTGGGAGAAGAACGTCCTTTCCGCTTCATTGAAGAAGAGAAGAAAGGCTTCCTCAAACGCTTGTTCG
GAGGATAAGGATCCGGCTGCTAACAAAGCCCGAAAGGAAGCTGAGTTGGCTGCTGCCACCGCTGAGC
AATAACTAGCATAACCCCTTGGGGCCTCTAAACGGGTCTTGAGGGGTTTTTTGATTGGGTATCGGATCC
CGGGCCCGTCGACTGCAGAGGCCTGCATGCAAGCTTGGCGTAATCATGGTCATAGCTGTTTCCTGTGT
GTAATACGACTCACTATAGGGGAATTGTGAGCGGATAACAATTCCCCTCTAGAAATAATTTTGTTTAACT
TTAAGAAGGAGATATACATATGGCGCTGCTGGATTTCTTTCTGAGCCGTAAGAAAAACACCGCGAACAT
CGCGAAAGAGCGTCTGCAAATCATTGTTGCGGAGCGTCGTCGTAGCGATGCGGAACCGCACTACCTG
CCGCAGCTGCGTAAAGATATCCTGGAAGTGATTTGCAAGTATGTTCAAATTGACCCGGAGATGGTGACC
GTTCAGCTGGAACAAAAGGACGGTGATATCAGCATTCTGGAGCTGAACGTTACCCTGCCGGAAGCGGA
GGAACTGAAGTAAGGATCCGGCTGCTAACAAAGCCCGAAAGGAAGCTGAGTTGGCTGCTGCCACCGC
TGAGCAATAACTAGCATAACCCCTTGGGGCCTCTAAACGGGTCTTGAGGGGTTTTTTG

#### DNA sequence of the *ftsA-minD-minE* construct

TAATACGACTCACTATAGGGGAATTGTGAGCGGATAACAATTCCCCTCTAGAAATAATTTTGTTTAACTT
TAAGAAGGAGATATACATATGATCAAGGCGACCGACCGTAAGCTGGTTGTGGGCCTGGAGATTGGCAC
CGCGAAGGTTGCGGCGCTGGTTGGCGAGGTTCTGCCGGATGGTATGGTTAACATTATCGGCGTTGGTA
GCTGCCCGAGCCGTGGCATGGACAAAGGTGGTGTGAACGACCTGGAAAGCGTGGTTAAGTGCGTGCA
GCGTGCGATTGACCAGGCGGAGCTGATGGCGGACTGCCAAATCAGCAGCGTTTACCTGGCGCTGAGC
GGCAAGCACATCAGCTGCCAAAACGAGATTGGTATGGTGCCGATTAGCGAAGAGGAAGTTACCCAGGA
AGATGTGGAGAACGTGGTTCACACCGCGAAAAGCGTTCGTGTGCGTGATGAACACCGTGTGCTGCACG
TTATCCCGCAAGAATACGCGATCGATTACCAGGAAGGTATCAAAAACCCGGTTGGTCTGAGCGGTGTT
CGTATGCAGGCGAAAGTGCACCTGATTACCTGCCACAACGATATGGCGAAGAACATTGTGAAAGCGGT
TGAACGTTGCGGTCTGAAGGTTGACCAGCTGATCTTCGCGGGTCTGGCGAGCAGCTACAGCGTTCTGA
CCGAAGATGAGCGTGAACTGGGTGTTTGCGTTGTGGATATCGGCGGTGGCACGATGGATATCGCGGT
GTATACCGGTGGCGCGCTGCGTCACACCAAAGTGATTCCGTATGCGGGTAACGTGGTTACCAGCGACA
TCGCGTACGCGTTTGGCACCCCGCCGAGCGATGCGGAGGCGATCAAAGTGCGTCACGGTTGCGCGCT
GGGTAGCATTGTGGGTAAAGATGAGAGCGTGGAAGTTCCGAGCGTTGGTGGCCGTCCGCCGCGTAGC

CTGCAACGTCAGACCCTGGCGGAAGTTATCGAGCCGCGTTACACCGAACTGCTGAACCTGGTGAACGA
AGAGATCCTGCAACTGCAAGAGAAACTGCGTCAGCAAGGTGTTAAGCACCACCTGGCGGCGGGCATT
GTTCTGACCGGCGGTGCGGCGCAGATCGAAGGTCTGGCGGCGTGCGCGCAACGTGTTTTCCACACCC
AAGTTCGTATCGGTGCGCCGCTGAACATCACCGGTCTGACCGATTACGCGCAAGAGCCGTACTATAGC
ACCGCGGTTGGTCTGCTGCACTATGGCAAAGAGAGCCACCTGAACGGCGAGGCGGAAGTGGAGAAGC
GTGTGACCGCGAGCGTTGGTAGCTGGATTAAGCGTCTGAATAGCTGGCTGCGTAAGGAGTTCTAAGGA
TCCGGCTGCTAACAAAGCCCGAAAGGAAGCTGAGTTGGCTGCTGCCACCGCTGAGCAATAACTAGCAT
AACCCCTTGGGGCCTCTAAACGGGTCTTGAGGGGTTTTTTGATCGGATCCCGGGCCCGTCGACTGCAG
AGGCCTGCATGCAAGCTTGGCGTAATCATGGTCATAGCTGTTTCCTGTGTGCTGCCCGCTTTCCAGTCG
GGAAACCTGTCGTGCCAGCTGCATTAATGAATCGGCCAACGCCAGTCACGACGTTGTAAAACGACGGC
CAGTGAATTCGAGCTCGGTACCTCGCGAATGCATCTAGATGCGAAATTAATACGACTCACTATAGGGGA
ATTGTGAGCGGATAACAATTCCCCTCTAGAAATAATTTTGTTTAACTTTAAGAAGGAGATATACATATGG
CACGCATTATTGTTGTTACTTCGGGCAAAGGGGGTGTTGGTAAGACAACCTCCAGCGCGGCCATCGCC
ACTGGTTTGGCCCAGAAGGGAAAGAAAACTGTCGTGATAGATTTTGATATCGGCCTGCGTAATCTCGAC
CTGATTATGGGTTGTGAACGCCGGGTCGTTTACGATTTCGTCAACGTCATTCAGGGCGATGCAACGCTA
AATCAGGCGTTAATTAAAGATAAGCGTACTGAAAATCTCTATATTCTGCCGGCATCGCAAACACGCGAT
AAAGATGCCCTCACCCGTGAAGGGGTCGCCAAAGTTCTTGATGATCTGAAAGCGATGGATTTTGAATTT
ATCGTTTGTGACTCCCCGGCAGGGATTGAAACCGGTGCGTTAATGGCACTCTATTTTGCAGACGAAGC
CATTATTACCACCAACCCGGAAGTCTCCTCAGTACGCGACTCTGACCGTATTTTAGGCATTCTGGCGTC
GAAATCACGCCGCGCAGAAAATGGCGAAGAGCCTATTAAAGAGCACCTGCTGTTAACGCGCTATAACC
CAGGCCGCGTAAGCAGAGGTGACATGCTGAGCATGGAAGATGTGCTGGAGATCCTGCGCATCAAACT
CGTCGGCGTGATCCCAGAGGATCAATCAGTATTGCGCGCCTCTAACCAGGGTGAACCGGTCATTCTCG
ACATTAACGCCGATGCGGGTAAAGCCTACGCAGATACCGTAGAACGTCTGTTGGGAGAAGAACGTCCT
TTCCGCTTCATTGAAGAAGAGAAGAAAGGCTTCCTCAAACGCTTGTTCGGAGGATAAGGATCCGGCTG
CTAACAAAGCCCGAAAGGAAGCTGAGTTGGCTGCTGCCACCGCTGAGCAATAACTAGCATAACCCCTT
GGGGCCTCTAAACGGGTCTTGAGGGGTTTTTTGATTGGGTATCGGATCCCGGGCCCGTCGACTGCAGA
GGCCTGCATGCAAGCTTGGCGTAATCATGGTCATAGCTGTTTCCTGTGTGTAATACGACTCACTATAGG
GGAATTGTGAGCGGATAACAATTCCCCTCTAGAAATAATTTTGTTTAACTTTAAGAAGGAGATATACATA
TGGCGCTGCTGGATTTCTTTCTGAGCCGTAAGAAAAACACCGCGAACATCGCGAAAGAGCGTCTGCAA
ATCATTGTTGCGGAGCGTCGTCGTAGCGATGCGGAACCGCACTACCTGCCGCAGCTGCGTAAAGATAT
CCTGGAAGTGATTTGCAAGTATGTTCAAATTGACCCGGAGATGGTGACCGTTCAGCTGGAACAAAAGG
ACGGTGATATCAGCATTCTGGAGCTGAACGTTACCCTGCCGGAAGCGGAGGAACTGAAGTAAGGATCC
GGCTGCTAACAAAGCCCGAAAGGAAGCTGAGTTGGCTGCTGCCACCGCTGAGCAATAACTAGCATAAC
CCCTTGGGGCCTCTAAACGGGTCTTGAGGGGTTTTTTG

**DNA sequence QconCAT**

TAATACGACTCACTATAGGGAGACCACAACGGTTTCCCTCTAGAAATAATTTTGTTTAACTTTAAGAAGG
AGATATACATATGCGGGGTTCTCATCATCATCATCATCATGGTATGGCTAGCATGACTGGTGGACAGCA
AATGGGTCGGGACCTGTACGACGACGACGATAAGAGCCTGTTTCCGAAAAGCAGCGTGGCGGTTAGC
TGGATTGGTAAAGCGGCGGCGGCGGGTGTGAGCGATGATGCGCGTATGGACCTGGCGAGCCAGTTTT
ACGAGAAGGCGGTGCTGGTTGATCTGGAACCGGGTGTTATTGCGCGTGCGACCAACCCGCTGGTGCG

TGCGGCGAGCCTGCTGGTTGACAAACTGGGCGAGAGCAGCAGCGGTAGCTATGTTCCGCGTAACATC
ATGGGTATTGAGGTGCCGGAAACCCTGGTTCACAAGACCATCTTCTTTAAACAGGCGAGCGAACAAAA
CTGGGCGAACTACAGCGCGGAGCAGAACCGTTTCGAAGGCGATACCCTGGTGAACCGTACCTACATCA
TCAGCATCCTGTTCAAGAACCAGATTGCGACCCTGGCGCAAAGCTATCGTGGTCTGGGTGCGGGTGCG
AACCCGGAAGTGGGTCGTGAATATCAAGACATTATCCGTATGCTGCAGCAAACCATCGAGCAGGCGCT
GCTGGAACAAGGTCGTCTGATTGTGCTGCTGCTGCTGGGCTTCGGCGAGCGTGCGTACGCGGATACC
GTTGAACGTGGCAGCAGCTTTACCCTGAGCGTGGTTCACCTGCACGAGGCGGAACCGAAGCTGCAGT
GGCAAGAGAGCGACGGTACCATCAAATACCGTCCGACCGACGATTTCGATGCGCGTTATACCGAACTG
CTGAACCTGGTGAACGAGGAAATTCTGCAGCTGCAAGAGAAGGACATCCTGGAAGTTATTTGCAAAGAT
GCGCTGAACCAGGCGGCGGACGATCTGAACCAACGTGTGGTTAACGACAACGCGCCGCAGACCGCGA
AAGATGCGCTGAGCCTGGCGCGTGGCGGTGTGAACGACCTGGAGAGCGTGGTTAAGATTATCGTGGT
TACCAGCGGTAAAGCGCTGGCGGGTGCGAGCGGTGACCGTGATACCATCGGCGACATTATCATTCTG
CCGCGTCTGAGCGATTACGGTGTTCAACTGCGTGCGCCGGTGGTTGTTCCGGCGGGTGTGGACGTGA
AAGCGATTAGCCTGAGCGTGCGTTAACTAGCATAACCCCTTGGGGCCTCTAAACGGGTCTTGAGGGGT
TTTTTG

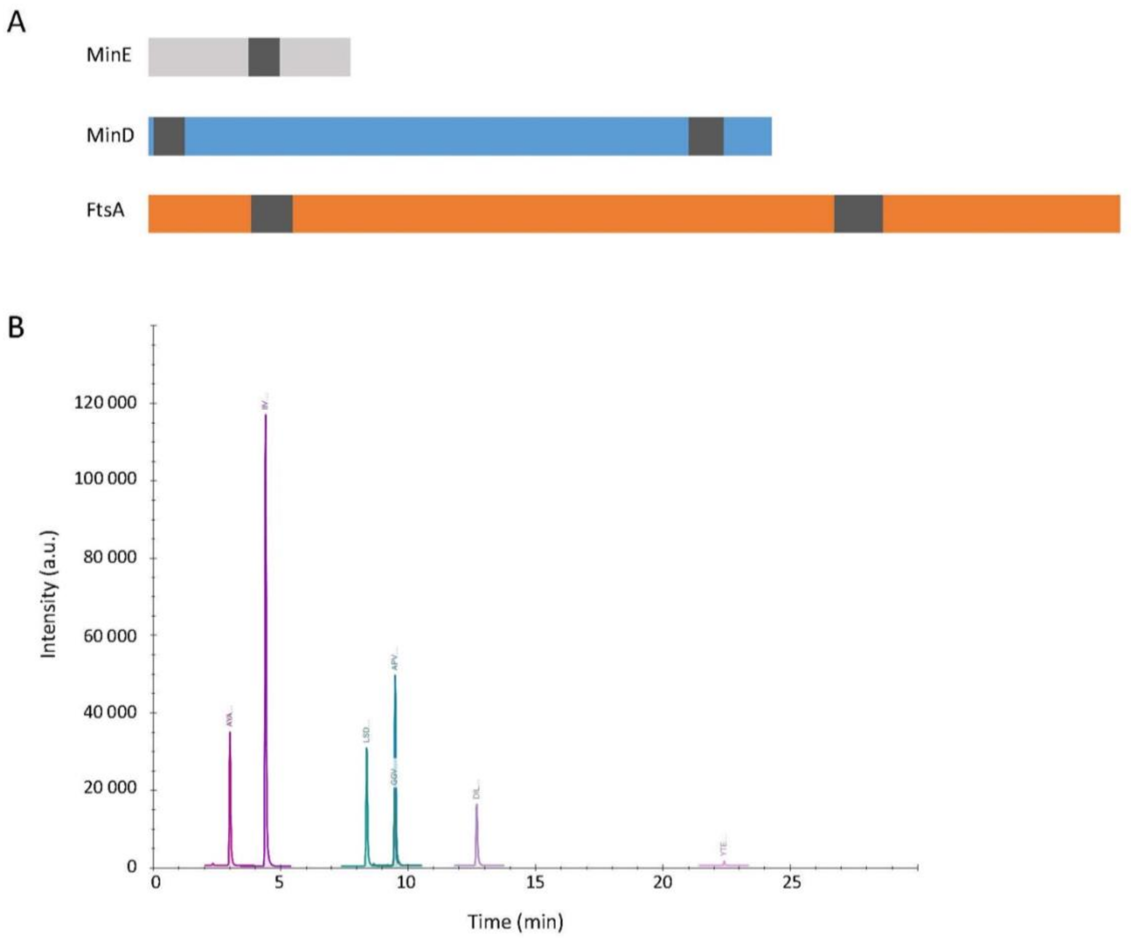


**Fig. S1.** Quantitative LC-MS analysis of cell-free synthesized FtsA, MinD and MinE in bulk reactions. (A) Cartoon depicting the position of the analyzed specific proteolytic peptides (dark grey domains) along the primary sequence of FtsA, MinD and MinE, from N-term (left) to C-term (right). (B) LC-MS chromatogram of the investigated peptides. From left to right: AYADTVER (MinD); IIVVTSGK (MinD); LSDYGVQLR (ribosomal protein S4); GGVNDLESVVK (FtsA); APVVVPAGVDVK (ribosomal protein L6); DILEVICK (MinE); YTELLNLVNEEILQLQEK (FtsA). The two peptides for ribosomal proteins were used for quality control.

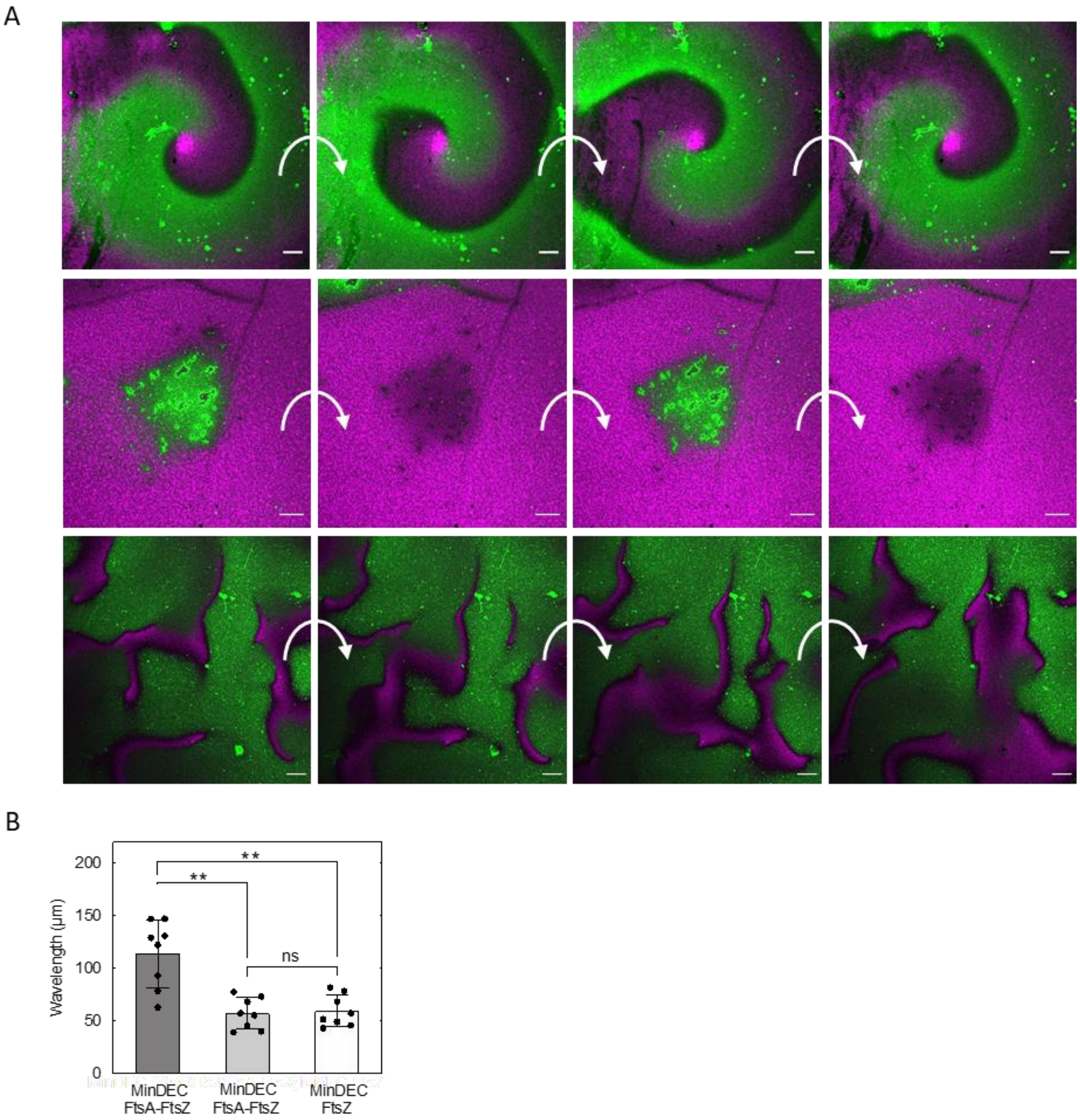


**Fig. S2**. MinDEC patterns and FtsA-FtsZ structures in supported membrane assays. (A) Composite fluorescence microscopy images of FtsA-FtsZ and MinDEC dynamic patterns on an SLB. Different patterns including spirals (top), standing (middle) and random (left) are shown. Arrows indicate time lapse between images. Experimental conditions and color coding are the same as in main text Fig. 2B and C. Scale bars are 10 µm. (B) Calculated wavelengths for MinDEC waves in areas where FtsA-FtsZ co-existed (dark grey), were excluded (light grey), and without expressed FtsA (white). Data are from three biological repeats and two to three fields of view have been analyzed per sample. Bar height represents the mean value and the error bar corresponds to the standard deviation. Symbols are values for individual fields of view aggregated from three biological replicates. Values obtained for different conditions were statistically compared by performing a two-tailed Welch's *t*-test. Asterisks indicate $P$ value <0.01, while 'ns' denotes a non-significant difference with $P$ value >0.05.

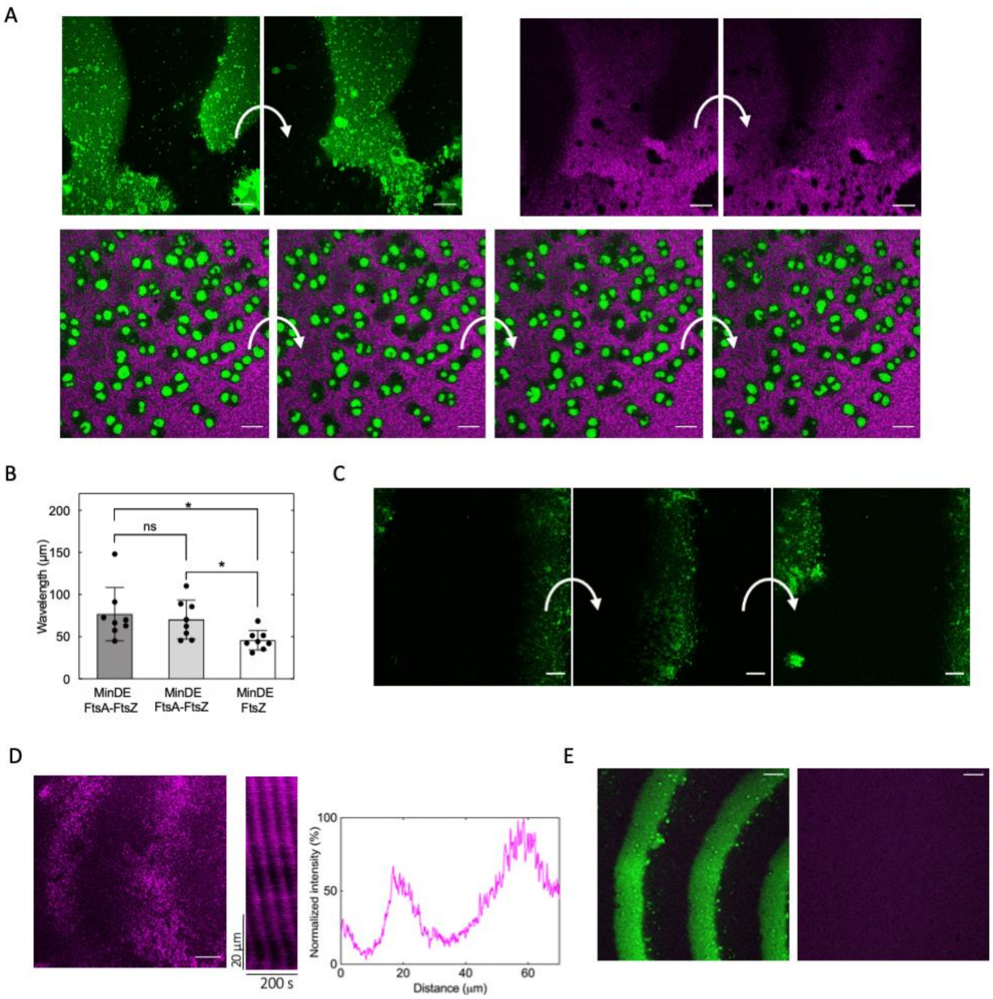


**Fig. S3.** Self-organized MinDE proteins, without MinC, can effectively rearrange FtsA-FtsZ filaments on supported membranes. (A) Fluorescence microscopy images showing that MinDE dynamic patterns rearrange the FtsA-FtsZ ring networks on an SLB. Examples of planar (top, split channels) and standing (bottom, merged channels) waves are shown. Arrows indicate time lapse between images. Experimental conditions and color coding are the same as in main text Fig. 3. Scale bars are 10 µm. (B) Calculated wavelengths for MinDE waves in areas where FtsA-FtsZ co-existed (dark grey), were excluded (light grey), and without expressed FtsA (white). Data are from three biological repeats and two to three fields of view have been analyzed per sample. Bar height represents the mean value and the error bar corresponds to the standard deviation. Symbols are values for individual fields of view aggregated from three biological replicates. Values obtained for different conditions were statistically compared by performing a two-tailed Welch's *t*-test. Asterisks indicate $P$ value <0.05, while 'ns' denotes a non-significant difference with $P$ value >0.05. (C) Fluorescence microscopy images of FtsA (0.4 µM FtsA-A488, green signal) dynamic patterns driven by MinDE on an SLB. Sharp propagating waves of FtsA are visible. Arrows indicate time lapse between images. (D) Fluorescence microscopy image of FtsA-FtsZ dynamic patterns on an SLB. Purified MinD reporter was omitted to rule out effects from a possible contamination with MinC. The corresponding kymograph is displayed, as well as the intensity profile of FtsZ signal along the direction of wave propagation. (E) Fluorescence microscopy images of FtsZ and MinDE dynamic patterns without expressed FtsA and purified eGFP-MinD (100 nM). In this condition, FtsZ is not recruited to the membrane (right image). Color coding: eGFP-MinD (green), FtsZ-A647 (magenta). Scale bars are 10 µm.

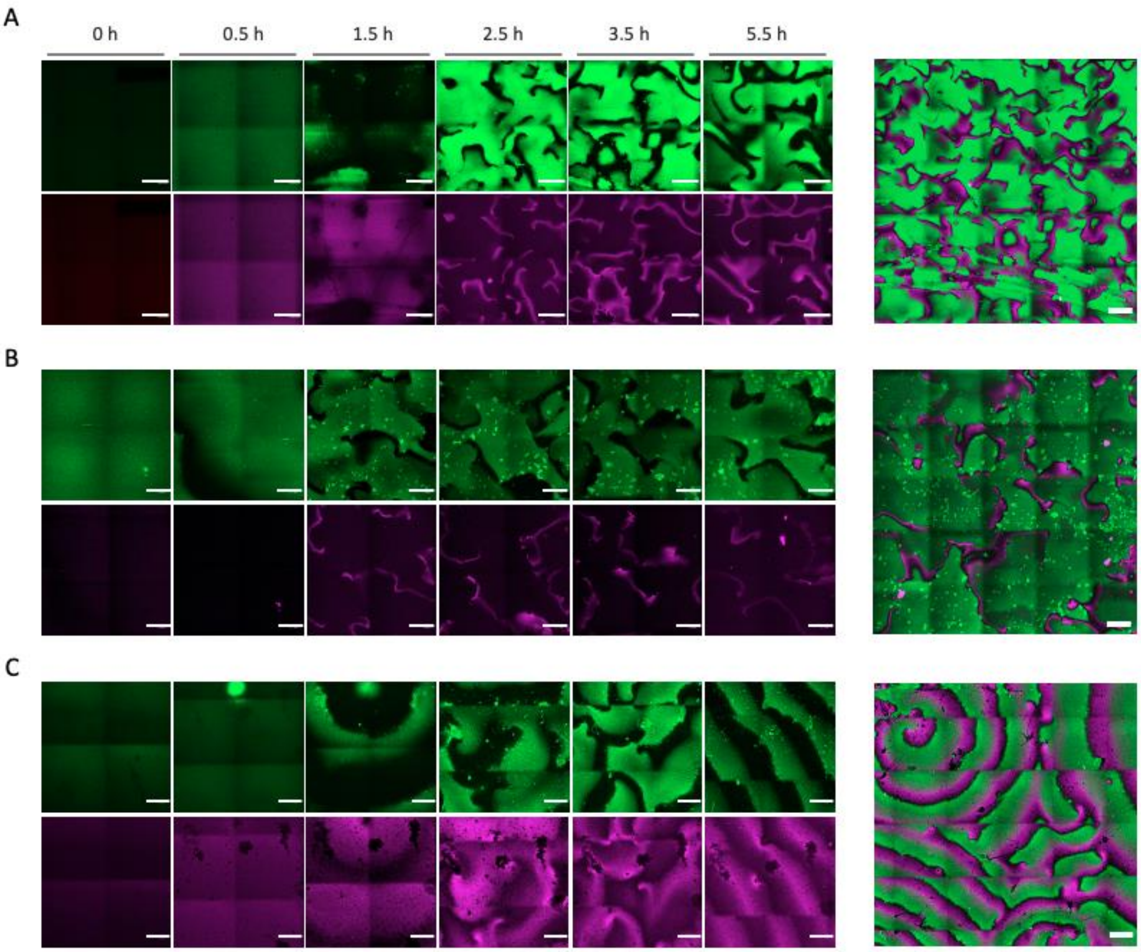


**Fig. S4.** Dynamic patterning of MinDE(C) and FtsA-FtsZ during in situ co-expression of FtsA, MinD and MinE. Samples were monitored up to 5.5 hours to study pattern creation and temporal evolution. (A) Two-by-two tile scan microscope images showing medium-scale organization of FtsA-FtsZ and MinDEC dynamic patterns at different time points. A mosaic image of 5×5 fields of view acquired after 3.5 hours of expression reveals large-scale, uniformly distributed waves of FtsA-FtsZ and MinDEC. Signals from eGFP-MinC and FtsZ-A647 are in green and magenta, respectively. The two channels were overlaid to compose the large mosaic image. Scale bars are 20 µm and 50 µm for the 2×2 and 5×5 tile scans, respectively. (B) Same as in A except that eGFP-MinC was replaced by a trace amount of purified eGFP-MinD (green). (C) Same as in B but in this sample the protein network self-organized into spirals and planar waves.

Dynamic FtsA-FtsZ patterns anti-correlating with the traveling MinDE(C) waves formed uniformly across the chamber. The coexistence of the two subsystems over the entire membrane surface contrasts with the spatial segregation of Min waves and FtsA-FtsZ filaments when pre-expressed proteins were added to the imaging chamber (main text Fig. 2 B and Fig.3 B). This different behavior may be attributed to protein-specific expression kinetics, which influences the coupling of the two subsystems at large length scales. For instance, MinE production being slow (main text Fig. 1 C) enables MinD and FtsA to bind homogenously all over the available membrane within the first hour until MinE concentration rises, triggering wave formation. Furthermore, in situ protein biogenesis may occur in close proximity to the lipid bilayer due to the intrinsic ability of ribosomes to bind to membranes, which potentially results in a different local concentration of FtsA and MinD comparing with pre-expressed proteins. In the presence of eGFP-MinC, rings of FtsA-FtsZ filaments formed after 1 hour of gene expression, concomitant to the emergence of MinDEC patterns that began in sporadic areas as standing waves (A). In the course of protein production, MinDEC self-organized into sharper traveling waves that colonized the whole chamber, causing stronger patterning of membrane-recruited FtsZ (A). In the assays conducted without MinC, eGFP-MinD was recruited to the membrane at the start of the reaction (B and C). After ~30 min, extensive dynamic patterns of MinDE appeared, followed by FtsZ surface waves that became more pronounced as more FtsA was synthesized. After 1 to 1.5 hours of gene expression, the anticorrelated concentration gradients of membrane-bound FtsA-FtsZ and MinDE became well defined. Here too, MinDE self-organized into crisper traveling waves as protein synthesis was progressing. Together, these results demonstrate that in situ cell-free gene expression can be exploited for timing protein-protein and protein-membrane interactions, which modulates the large-scale organization of the coupled Min-FtsA-FtsZ system.

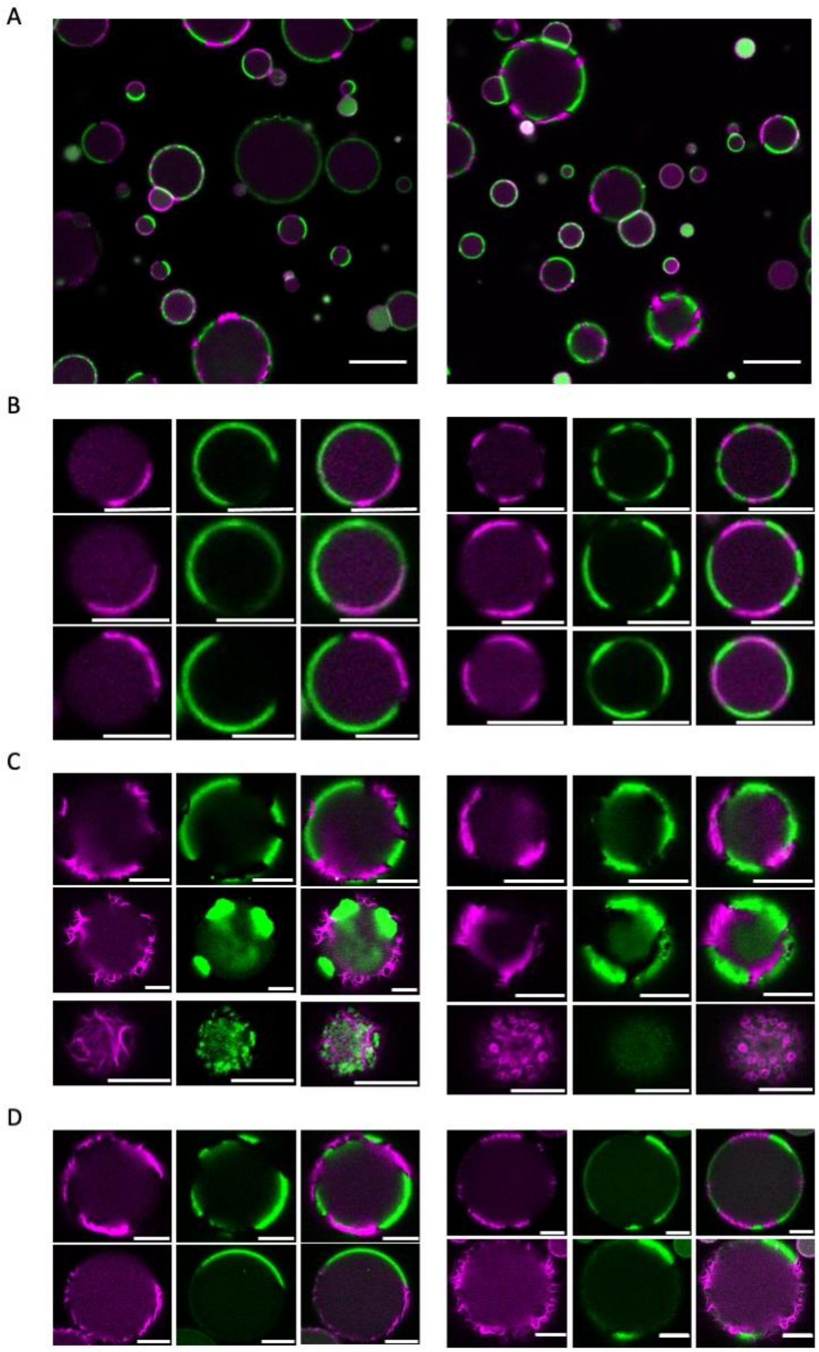


**Fig. S5.** Reconstitution of MinDEC and FtsA-FtsZ subsystems in water-in-oil droplets. Composite fluorescence microscopy images of droplet populations and a library of single droplet images with reconstituted MinDEC patterns and FtsA-FtsZ subsystems are shown. (A) Composite image of droplet population. Experimental conditions were as described in main text Fig. 4B. (B) Fluorescence images of droplets exhibiting antiphase dynamic patterns of MinDEC and FtsA-FtsZ. Experimental conditions were as described in main text Fig. 4B. (C) Same as in A, but images were acquired closer to the dome of the droplets. Multiple interfacial FtsZ polarization sites rearranged by the Min dynamics are visible. Experimental conditions were as described in main text Fig. 4B. (D) Same as in B-C, except that bigger droplets (diameter >20 µm) were imaged. Here, FtsZ cytoskeletal structures are clearly resolved. (A-D) Experimental conditions were as described in main text Fig. 4B. Color coding: eGFP-MinC (green), FtsZ-A647 (magenta). Scale bars are 10 µm.

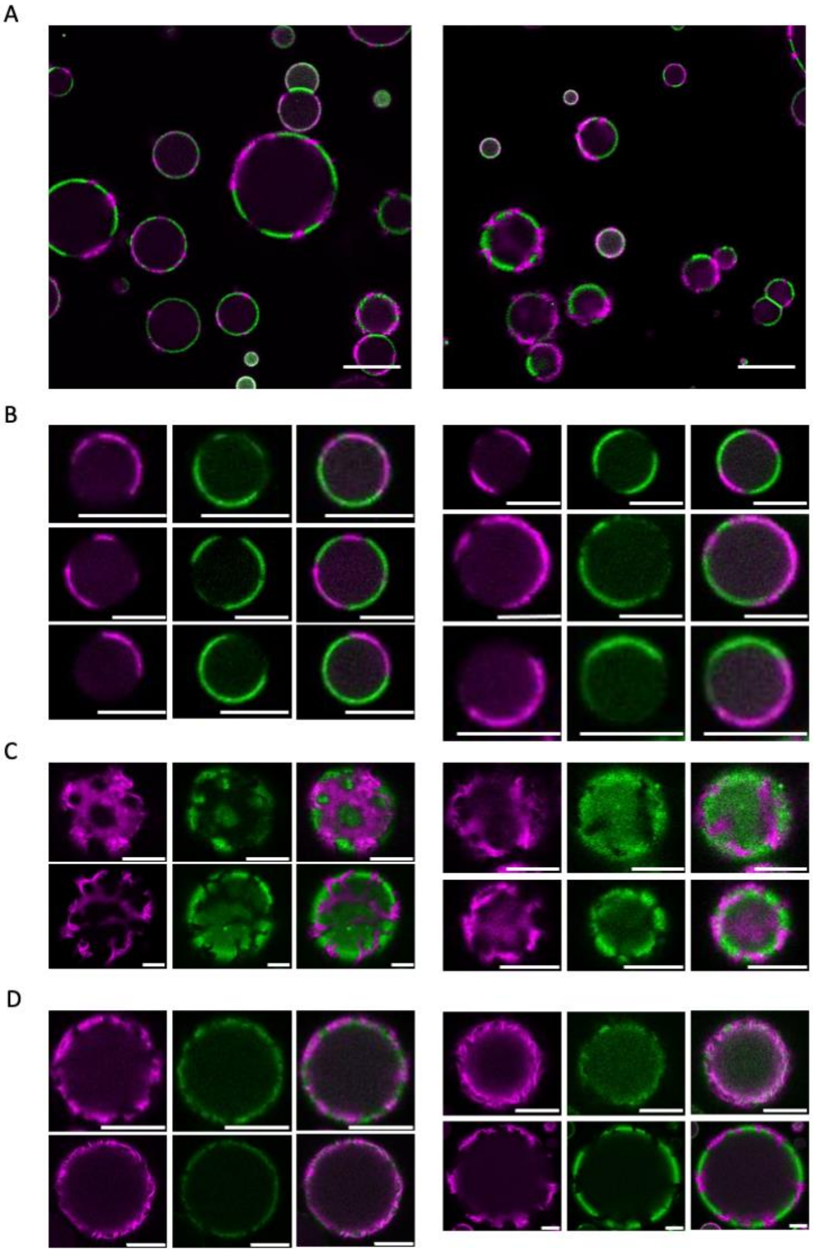


**Fig. S6.** Reconstitution of MinDE and FtsA-FtsZ subsystems in water-in-oil droplets. Composite fluorescence microscopy images of droplet populations and a library of single droplet images with reconstituted MinDE patterns and FtsA-FtsZ subsystems are shown. (A) Composite image of droplet population. Experimental conditions were as described in main text Fig. 4C. (B) Fluorescence images of droplets exhibiting antiphase dynamic patterns of MinDE and FtsA-FtsZ. Experimental conditions were as described in main text Fig. 4C. (C) Same as in A, but images were acquired closer to the dome of the droplets. Multiple interfacial FtsZ polarization sites rearranged by the Min dynamics are visible. Experimental conditions were as described in main text Fig. 4C. (D) Same as in B-C, except that bigger droplets (diameter >20 µm) were imaged. Here, FtsZ cytoskeletal structures are clearly resolved. Experimental conditions were as described in main text Fig. 4C. (A-D) Color coding: eGFP-MinD (green), FtsZ-A647 (magenta). Scale bars are 10 µm.

**Table S1.** List of primers used in this study

| Name | Sequence (5' to 3') |
|---|---|
| 173 ChD | CACACAGGAAACAGCTATGAC |
| 174 ChD | GAGTCAGTGAGCGAGGAAG |
| 365 ChD | CAGTCACGACGTTGTAAAACGAC |
| 420 ChD | GGAGAGGCGGTTTGCGTAT |
| 504 ChD | CTGCCCGCTTTCCAGTCGGGAAA |
| 1139 ChD | CATGGTCATAGCTGTTTCCTGTGTGTAATACGACTCACTATAGG |
| 1140 ChD | GGTTTCCCGACTGGAAAGCGGGCAGCAAAAAACCCCTCAAGACCCG |
| 1145 ChD | CCGTCGTTTTACAACGTCGTGACTGGCGTTGGCCGATTCATTAATGC |
| 1187 ChD | GGCGATTAAGTTGGGTAACG |

**Table S2.** Transitions of the MS/MS measurements for the proteolytic peptides of the indicated proteins. Accelerator voltage was kept constant at 4 eV.

| **Protein** | **Compound name** | **Precursor ion (m/z)** | **Product ion (m/z)** | **Collision energy (eV)** | **Ion name** |
|---|---|---|---|---|---|
| MinD | AYADTVER | 462.725 | 690.342 | 15.3 | y6 |
| MinD | AYADTVER | 462.725 | 619.305 | 15.3 | y5 |
| MinD | AYADTVER | 462.725 | 304.162 | 15.3 | y2 |
| MinD | AYADTVER | 462.725 | 235.108 | 15.3 | b2 |
| MinD | AYADTVER | 462.725 | 306.145 | 15.3 | b3 |
| MinD | AYADTVER | 462.725 | 621.288 | 15.3 | b6 |
| QconCAT | AYADTVER | 468.208 | 699.315 | 15.3 | y6 |
| QconCAT | AYADTVER | 468.208 | 627.281 | 15.3 | y5 |
| QconCAT | AYADTVER | 468.208 | 309.147 | 15.3 | y2 |
| QconCAT | AYADTVER | 468.208 | 237.102 | 15.3 | b2 |
| QconCAT | AYADTVER | 468.208 | 309.136 | 15.3 | b3 |
| QconCAT | AYADTVER | 468.208 | 627.270 | 15.3 | b6 |
| FtsA | YTELLNLVNEEILQLQEK | 1095.089 | 1243.653 | 34.9 | y10 |
| FtsA | YTELLNLVNEEILQLQEK | 1095.089 | 871.525 | 34.9 | y7 |
| FtsA | YTELLNLVNEEILQLQEK | 1095.089 | 758.441 | 34.9 | y6 |
| FtsA | YTELLNLVNEEILQLQEK | 1095.089 | 645.357 | 34.9 | y5 |
| FtsA | YTELLNLVNEEILQLQEK | 1095.089 | 517.298 | 34.9 | y4 |
| FtsA | YTELLNLVNEEILQLQEK | 1095.089 | 847.456 | 34.9 | b7 |
| QconCAT | YTELLNLVNEEILQLQEK | 1106.555 | 1257.611 | 34.9 | y10 |
| QconCAT | YTELLNLVNEEILQLQEK | 1106.555 | 881.495 | 34.9 | y7 |
| QconCAT | YTELLNLVNEEILQLQEK | 1106.555 | 767.414 | 34.9 | y6 |
| QconCAT | YTELLNLVNEEILQLQEK | 1106.555 | 653.333 | 34.9 | y5 |
| QconCAT | YTELLNLVNEEILQLQEK | 1106.555 | 523.280 | 34.9 | y4 |
| QconCAT | YTELLNLVNEEILQLQEK | 1106.555 | 855.432 | 34.9 | b7 |
| MinE | DILEVIC[+57]K | 495.270 | 874.507 | 15.6 | y7 |
| MinE | DILEVIC[+57]K | 495.270 | 761.423 | 15.6 | y6 |
| MinE | DILEVIC[+57]K | 495.270 | 648.339 | 15.6 | y5 |
| MinE | DILEVIC[+57]K | 495.270 | 471.245 | 15.6 | b4 |
| MinE | DILEVIC[+57]K | 495.270 | 570.313 | 15.6 | b5 |
| MinE | DILEVIC[+57]K | 495.270 | 683.397 | 15.6 | b6 |
| QconCAT | DILEVIC[+57]K | 499.757 | 882.483 | 15.6 | y7 |
| QconCAT | DILEVIC[+57]K | 499.757 | 768.402 | 15.6 | y6 |
| QconCAT | DILEVIC[+57]K | 499.757 | 654.321 | 15.6 | y5 |
| QconCAT | DILEVIC[+57]K | 499.757 | 475.233 | 15.6 | b4 |
| QconCAT | DILEVIC[+57]K | 499.757 | 575.299 | 15.6 | b5 |
| QconCAT | DILEVIC[+57]K | 499.757 | 689.380 | 15.6 | b6 |
| FtsA | GGVNDLESVVK | 558.798 | 903.478 | 18.3 | y8 |
| FtsA | GGVNDLESVVK | 558.798 | 789.435 | 18.3 | y7 |
| FtsA | GGVNDLESVVK | 558.798 | 674.408 | 18.3 | y6 |

**Table S2. *Continued***

| Protein | Compound name | Precursor ion (m/z) | Product ion (m/z) | Collision energy (eV) | Ion name |
|---|---|---|---|---|---|
| FtsA | GGVNDLESVVK | 558.798 | 561.324 | 18.3 | y5 |
| FtsA | GGVNDLESVVK | 558.798 | 432.282 | 18.3 | y4 |
| FtsA | GGVNDLESVVK | 558.798 | 214.119 | 18.3 | b3 |
| QconCAT | GGVNDLESVVK | 565.279 | 913.449 | 18.3 | y8 |
| QconCAT | GGVNDLESVVK | 565.279 | 797.412 | 18.3 | y7 |
| QconCAT | GGVNDLESVVK | 565.279 | 681.388 | 18.3 | y6 |
| QconCAT | GGVNDLESVVK | 565.279 | 567.306 | 18.3 | y5 |
| QconCAT | GGVNDLESVVK | 565.279 | 437.267 | 18.3 | y4 |
| QconCAT | GGVNDLESVVK | 565.279 | 217.110 | 18.3 | b3 |
| MinD | IIVVTSGK | 408.763 | 703.435 | 13.7 | y7 |
| MinD | IIVVTSGK | 408.763 | 590.351 | 13.7 | y6 |
| MinD | IIVVTSGK | 408.763 | 491.282 | 13.7 | y5 |
| QconCAT | IIVVTSGK | 413.250 | 711.411 | 13.7 | y7 |
| QconCAT | IIVVTSGK | 413.250 | 597.330 | 13.7 | y6 |
| QconCAT | IIVVTSGK | 413.250 | 497.265 | 13.7 | y5 |
| QconCAT | IIVVTSGK | 413.250 | 429.300 | 13.7 | b4 |
| QconCAT | IIVVTSGK | 413.250 | 531.345 | 13.7 | b5 |
| Ribosomal protein S4 | LSDYGVQLR | 525.783 | 850.442 | 17.3 | y7 |
| Ribosomal protein S4 | LSDYGVQLR | 525.783 | 735.415 | 17.3 | y6 |
| Ribosomal protein S4 | LSDYGVQLR | 525.783 | 572.351 | 17.3 | y5 |
| Ribosomal protein S4 | LSDYGVQLR | 525.783 | 635.304 | 17.3 | b6 |
| QconCAT | LSDYGVQLR | 532.263 | 949.438 | 17.3 | y8 |
| QconCAT | LSDYGVQLR | 532.263 | 861.409 | 17.3 | y7 |
| QconCAT | LSDYGVQLR | 532.263 | 745.385 | 17.3 | y6 |
| QconCAT | LSDYGVQLR | 532.263 | 581.325 | 17.3 | y5 |
| QconCAT | LSDYGVQLR | 532.263 | 541.220 | 17.3 | b5 |
| QconCAT | LSDYGVQLR | 532.263 | 641.286 | 17.3 | b6 |
| Ribosomal protein L6 | APVVVPAGVDVK | 575.845 | 883.525 | 18.9 | y9 |
| Ribosomal protein L6 | APVVVPAGVDVK | 575.845 | 784.456 | 18.9 | y8 |
| Ribosomal protein L6 | APVVVPAGVDVK | 575.845 | 685.388 | 18.9 | y7 |
| Ribosomal protein L6 | APVVVPAGVDVK | 575.845 | 268.166 | 18.9 | b3 |
| Ribosomal protein L6 | APVVVPAGVDVK | 575.845 | 367.234 | 18.9 | b4 |
| Ribosomal protein L6 | APVVVPAGVDVK | 575.845 | 466.302 | 18.9 | b5 |
| QconCAT | APVVVPAGVDVK | 582.326 | 893.495 | 18.9 | y9 |
| QconCAT | APVVVPAGVDVK | 582.326 | 793.430 | 18.9 | y8 |
| QconCAT | APVVVPAGVDVK | 582.326 | 693.364 | 18.9 | y7 |
| QconCAT | APVVVPAGVDVK | 582.326 | 271.157 | 18.9 | b3 |
| QconCAT | APVVVPAGVDVK | 582.326 | 371.222 | 18.9 | b4 |
| QconCAT | APVVVPAGVDVK | 582.326 | 471.288 | 18.9 | b5 |

**Movie S1 (separate file).** Video of FtsA-FtsZ and MinDEC dynamic patterns on a supported membrane. The experimental conditions are identical as in Fig. 2C. Colors: eGFP-MinC (green), FtsZ-A647 (magenta). Scale bar is 10 µm.

**Movie S2 (separate file).** Video of FtsZ and MinDEC dynamic patterns without expressed FtsA on an SLB. In the absence of FtsA, FtsZ colocalizes with MinDEC waves. The experimental conditions are identical as in Fig. 2E. Colors: eGFP-MinC (green), FtsZ-A647 (magenta). Scale bar is 10 µm.

**Movie S3 (separate file).** Video of FtsA-FtsZ and MinDE dynamic patterns on an SLB. MinDE proteins, without MinC, regulate FtsA-FtsZ spatial organization. The experimental conditions are identical as in Fig. 3. Colors: eGFP-MinD (green), FtsZ-A647 (magenta). Scale bar is 10 µm.

**Movie S4 (separate file).** Videos of FtsA-FtsZ and MinDE dynamic patterns on an SLB for areas showing distinct rings and filaments of FtsZ. Waves effectively rearranging FtsA-FtsZ filaments and low-amplitude propagating waves of FtsZ anticorrelating with MinDE patterns are played one after the other. The experimental conditions are identical as in Fig. 3. Colors: eGFP-MinD (green), FtsZ-A647 (magenta). Scale bars are 10 µm.

**Movie S5 (separate file).** Videos of MinDE(C) and FtsA-FtsZ exhibiting antiphase dynamic patterns in water-in-oil droplets. One field of view is shown. The experimental conditions are identical as in Fig. 5. Colors: eGFP-MinC/D (green), FtsZ-A647 (magenta). Scale bars are 10 µm.

**Movie S6 (separate file).** Videos of two droplets exhibiting antiphase dynamic patterns of either MinDEC and FtsA-FtsZ (corresponding to the droplets in Fig. 5B), or MinDE and FtsZ (corresponding to the droplets in Fig. 5C), are played one after the other. The experimental conditions are identical as in Fig. 5. Colors: eGFP-MinC/D (green), FtsZ-A647 (magenta). Scale bars are 10 µm.

**Movie S7 (separate file).** Videos of droplets exhibiting antiphase dynamic patterns of MinDE(C) and FtsA-FtsZ (corresponding to the droplets in Fig. 5D, exact condition is as specified) are played side by side. The two droplets have a diameter >20 µm, allowing visualization of FtsZ cytoskeletal structures extending to the lumen. The experimental conditions are identical as in Fig. 5. Colors: eGFP-MinC/D (green), FtsZ-A647 (magenta). Scale bars are 10 µm.
